# Coastal winds and larval fish abundance indicate a recruitment mechanism for southeast Australian estuarine fisheries

**DOI:** 10.1101/2020.06.24.170068

**Authors:** Hayden T. Schilling, Charles Hinchliffe, Jonathan P. Gillson, Anthony Miskiewicz, Iain M. Suthers

**Affiliations:** Sydney Institute of Marine Science, Chowder Bay Road, Mosman, 2088, Australia; Centre for Marine Science & Innovation, UNSW Australia, 2052 Australia; The Centre for Environment, Fisheries and Aquaculture Science, Pakefield Road, Lowestoft, NR33 0HT, United Kingdom; Australian Museum, College St., Sydney 2000, Australia

**Keywords:** larval fish, retention, wind driven upwelling, fisheries production, Harald Dannevig, downwelling, estuarine recruitment

## Abstract

Coastal winds transport water masses and larval fish onshore or offshore which may influence estuarine recruitment, yet our understanding of the mechanism underlying this relationship is limited. Here, we combine datasets from a historical database of larval fish off southeast Australia with a high-resolution atmospheric reanalysis model to show that normalised abundance of coastally spawned larvae increased with weak to moderate upwelling favourable winds 14 days prior to sampling. The increase in abundance may reflect increased nutrient and plankton availability for larval fish. Normalised larval abundance decreased following strong upwelling favourable winds but increased after onshore (downwelling favourable) winds, due to wind-driven transport. By combining a commercial estuarine fisheries catch-rate dataset (4 species, 8 estuaries, 10 years) and the high-resolution atmospheric reanalysis model, we show that negative effects of upwelling favourable winds during the spawning period can be detected in lagged estuarine commercial fisheries catch rates (lagged by 2 – 8 years depending on species’ growth rates), potentially representing the same mechanism proposed for larval fish. Upwelling favourable winds in the southeast Australian region have increased since 1850 while onshore winds have decreased, which may reduce larval recruitment to estuaries. Coastal winds are likely an important factor for estuarine recruitment in the southeast Australian region and future research on the estuarine recruitment of fish should incorporate coastal winds.

## Introduction

Strong year classes of marine fish have long interested fisheries oceanographers, with many studies attempting to better understand the drivers of variability in population dynamics (Houde, 2008). Previous studies have linked strong year classes with abiotic and biotic factors during the larval period, with hypotheses either focusing on increased growth or reduced mortality resulting in a higher recruitment (Garvine *et al*., 1997; Helbig and Pepin, 1998; Pepin, 2016). Some of the key factors influencing recruitment include the biomass of the spawning stock, location and timing of spawning, the vertical distribution of eggs and larvae and upwelling generated productivity. The spawning stock biomass (SSB) is the basis for recruitment success (Cury *et al*., 2014) and in combination with fecundity, egg viability and sex ratio, determines the number of eggs and larvae which subsequent stochastic processes act upon to shape recruitment (Kell *et al*., 2016).

Stochastic processes are often more influential than SSB due to the high variability in mortality and subsequent recruitment (Szuwalski *et al*., 2015). The location and timing of spawning events is also important for overall recruitment, with seasonal and interannual variations in oceanography often shaping dispersal and potential recruitment (Ciannelli *et al*., 2015; Schilling *et al*., 2020). As larvae have limited swimming potential in their early stages (hence oceanographic driven dispersal), larvae which can control their vertical positioning often exploit vertical variation in cross-shelf flow on continental shelves to remain near favourable areas for juvenile habitat, potentially increasing recruitment (Hare and Govoni, 2005; Ospina-Alvarez *et al*., 2018).

Productivity, which is often generated by upwelling (Everett *et al*., 2014), is often suggested as a driver of faster growth in fish larvae through the intermediate production of phytoplankton and zooplankton thereby reducing mortality from predation, as the larvae increase in swimming ability (Anderson, 1988; McFarlane and Beamish, 1992), and increasing recruitment (Bailey and Houde, 1989). On the other hand, upwelling can also have negative effects in some circumstances due to offshore transport away from suitable recruitment habitat. While the influence of upwelling has been clearly demonstrated in some regions including the southern Benguela upwelling region (Wilhelm *et al*., 2005) and the Northeast Atlantic (Santos *et al*., 2004), this process is likely region specific and remains uncertain in many regions, including the southwest Pacific.

The coastal waters of eastern Australia are dominated by the East Australian Current (EAC; Oke *et al*., 2019). The EAC generally transports larvae poleward along a narrow continental shelf (<30km) to coastal and estuarine nursery habitats (Roughan *et al*., 2011; Schilling *et al*., 2020). Together with the EAC, onshore winds are an important driver of upwelling and downwelling through Ekman transport mechanisms (Roughan and Middleton, 2002; Schaeffer *et al*., 2013, 2014). Wind is an important driver of cross-shelf flows in the area (McClean-Padman and Padman, 1991; Middleton *et al*., 1996). Winds from the northeast (NE) blow along the coastline, promoting offshore Ekman transport and upwelling of cold nutrient-rich water along the coast, while winds from the southeast (SE), promoting onshore transport and downwelling (Griffin and Middleton, 1992; Middleton *et al*., 1996).

It was first hypothesised that onshore winds may be affecting nearshore retention and potentially driving fluctuations in commercial catches in southeastern Australian estuaries in the early 20^th^ century (Dannevig, 1907; Suthers *et al*., 2020). Dannevig (1907) hypothesised that onshore winds reduced unfavourable advection of fish eggs and larvae, retaining them nearer the coast and therefore increasing recruitment to estuarine fisheries. When testing this hypothesis, Dannevig (1907) found a positive correlation between estuarine commercial catch rates and onshore winds lagged by three or four years (Suthers *et al*., 2020). Since the inception of this hypothesis, many studies have shown relationships between wind and juvenile recruitment with the effects largely being system specific (Nelson *et al*., 1977; Caputi *et al*., 2001; Takeshige *et al*., 2013; Wilson and Laman, 2021), likely due to complex estuary specific recruitment processes (Boehlert and Mundy, 1988).

The winds in many of the world’s coastal upwelling systems are being altered due to climate change, resulting in both increases (California, Benguela and Humboldt systems) and decreases (e.g. Iberian system) in upwelling favourable winds (Sydeman *et al*., 2014). Changing winds are likely to have a range of effects including enrichment of waters, regional changes in stratification and basin-scale changes in thermocline structure, all of which may influence the productivity of local fisheries (Bakun *et al*., 2010). Concerns have been expressed with regards to changing coastal winds altering upwelling regimes which provide important nutrients to ecosystems (Bakun and Weeks, 2008; Bakun *et al*., 2010; Sorte, 2013). In addition to changes in nutrient supply, variation in upwelling patterns may also affect the advection and recruitment of larval fish to estuaries.

Estuary specific characteristics also contribute to variation in commercial catch rates through both short term catchability effects and longer term productivity differences among estuaries. Within estuarine environments, fish and fisher behaviour can change in response to external effects such as weather or freshwater flow (Gillson, 2011). Key estuarine fisheries families such as Sparidae have been shown to reverse their diel behaviour patterns following heavy rain (Payne *et al*., 2013), with the catch rates of many species differing between drought and non-drought conditions (Gillson *et al*., 2009). Within eastern Australia, it has been demonstrated that there are consistent differences in hydrology and fisheries production between types of estuaries defined by their geomorphology (Pease, 1999; Roy *et al*., 2001). When investigating variation in estuarine fisheries production, potentially due to larval recruitment effects, it is therefore necessary to control for differences originating from differences in freshwater flow or estuarine type.

In the present study, we test the original hypothesis of Dannevig (1907) using both larval fish abundance and commercial fisheries data. To evaluate offshore and onshore winds (upwelling favourable and downwelling favourable respectively) as a driver of estuarine fisheries production in southeast Australia, model-based estimates of surface wind were first compared to normalised abundances of 132 taxa of coastally spawned fish larvae (1990 - 2016), and then to estuarine fishery catches (4 species, 8 estuaries, July 1997 – June 2007). Our present study extends the idea of Dannevig (1907) by investigating both larval fish and commercial catch data to test three specific aims. Firstly, we examine whether coastal winds influence the abundance of coastally spawned larval fish near the coast. Secondly, we evaluate whether the same winds during the spawning period influence commercial fisheries harvest when lagged by an appropriate growth period for each species. Finally, we explore changes in the upwelling and downwelling (onshore) favourable winds since 1850. We expect that coastally spawned larval abundance will be greater during periods of onshore winds. If the onshore transport of coastal fish larvae does occur during downwelling favourable winds, then the effects of this retention should result in increased larval supply to estuaries, and assuming juvenile fish are resident in estuaries until size of capture, may result in a detectable effect on commercial fisheries catch rates.

## Method

### Dataset Descriptions

#### Wind Data

To provide a consistent estimate of winds, we used the wind speed and direction data from the Australian Bureau of Meteorology Atmospheric high-resolution Regional Reanalysis for Australia (BARRA; Su *et al*., 2019, 2020). This re-analysis product provides hourly wind speeds at a 12 km resolution over the Australian domain with downscaled 1.5 km resolution within several sub-domains between 1990 and 2019. For all analyses in the present study, we used near-surface (10 m) wind data from the 1.5 km resolution data from within the Eastern New South Wales subdomain (Su *et al*., 2020). This 1.5 km reanalysis model demonstrates high agreement with observations, with higher skill than the 12 km, particularly in coastal areas (Su *et al*., 2020).

To provide a long-term context, we also used wind speed and directions from the 20^th^ Century Reanalysis V2c data collated by the NOAA/OAR/ESRL PSD (www.esrl.noaa.gov/psd/), which provides three-hourly wind speed and direction data from 1850 to 2014 at a resolution of ≈200 km (Compo *et al*., 2015). As both wind products are reanalysis datasets, the models are constrained by observed values.

#### Larval Fish Data

To investigate the response of larval fish abundance to wind, we used data from the Australian Integrated Marine Observing System (IMOS) Larval Fish Database (Smith *et al*., 2018). Fish larvae were all collected from the top 150m of water using horizontal and oblique tows from variety of plankton nets with a 300 – 500 μm mesh size (see Smith *et al*. (2018) for full dataset description and Table S1 for a summary of the samples included in the current analysis). Each sample refers to a single plankton tow and the abundance of each taxon collected. While the database contains abundance information for 218 taxa collected over 12 research projects (1983 - 2016; Smith et al., 2018), the present study selected only seven of these projects that matched the temporal resolution of the wind data (1990 – 2016; Table S1) for analysis.

#### Commercial Fisheries Data

To assess whether the effects of onshore winds during the spawning period, on the abundance of coastally spawned fish larvae could be detected at the commercial fisheries scale, we used an estuarine catch-per-unit-effort (CPUE) dataset from eastern Australia (Gillson *et al*., 2009). This dataset consisted of ten years of monthly CPUE data (combined to annual values; July 1997 – June 2007) from gillnet fisheries in located inside eight estuaries along the east Australian coast (Roy *et al*., 2001; Table S1). An extensive description of the dataset is given in Gillson *et al*. (2009). Briefly, these CPUE data were derived from monthly catch (kg of harvest) and effort (fishing days) for four fish species; yellowfin bream (*Acanthopagrus australis*; Sparidae), sea mullet (*Mugil cephalus*; Mugilidae), dusky flathead (*Platycephalus fuscus*; Platycephalidae) and sand whiting (*Sillago ciliata*; Sillaginidae). These fish species represent the dominant contribution to both commercial and recreational estuarine fisheries harvest. All four of these species are known to spawn in nearshore coastal waters before juveniles recruit to estuaries (spawning seasons and references detailed in Table 1). Sand whiting and dusky flathead larvae are usually found at deeper depths (15-50 m) while sea mullet larvae are found in surface waters (Collins and Stender, 1989). Yellowfin bream larvae are found equally throughout the top 30 m of the water column, with some larvae found up to 70 m deep (Gray, 1993; Gray *et al*., 2019). Luderick (*Girella tricuspidata*; Girellidae) was in the original data (Gillson *et al*., 2009), but was not included in the present study due to the variable spawning periods between populations in this region (Gray *et al*., 2012). While CPUE is often not considered a robust measure of fish abundance (Richards and Schnute, 1986; Harley *et al*., 2001; Haggarty and King, 2006), for our purpose gillnet CPUE provided a measure of fisheries catch (≥80mm mesh size) using a passive gear type to assess the impacts of variation in recruitment on catch rates and presented the most consistent method available from which an index of abundance could be inferred (Gillson *et al*., 2009). Due to reporting requirements at the time, effort represents total gillnet effort on monthly timescale, undifferentiated by species. There is some seasonality to the fishing effort which is generally higher in winter months, corresponding to increased CPUE for flathead and bream, suggesting fishers may target bream and flathead in winter months. As we analysed our data on an annual time scale and there are no changes in annual effort (Figure S3), the seasonality is consistent between years.

**Table 1.** Details of the spawning season and modal age of capture in gillnet fisheries for four coastally spawning fish species, used to define the choice of lags for winds during the spawning season with an example of the wind intervals used for the annual catch-per-unit-effort (CPUE) between July 1997 and June 1998. The ranges of harvested ages (> 5 % of the catch) is shown in brackets with the modal harvested age. Full age distributions can be seen in Gray *et al*. (2015).

| Species | Spawning season | Modal harvested age<br>(Range; Gray *et al.*, 2015) | Example wind intervals used for annual CPUE for July 1997 – June 1998 |
| --- | --- | --- | --- |
| bream<br>(*Acanthopagrus australis*) | October – February (Ochwada-Doyle *et al.*, 2012) | 7 (4 – 11) | October 1989 – February 1990 |
| dusky flathead<br>(*Platycephalus fuscus*) | December – March (Gray and Barnes, 2015) | 3 (2-5) | December 1993 – March 1994 |
| sand whiting<br>(*Sillago ciliata*) | December – February (Burchmore *et al.*, 1988) | 6 (3 -10) | December 1991 – February 1992 |
| sea mullet<br>(*Mugil cephalus*) | April – July (Stewart *et al.*, 2018) | 2 (1-5) | April – July 1995 |

### Data Preparation

All data preparation and analysis was conducted using R v4.0.2 (R Core Team, 2020).

#### Wind Data

Wind speed and direction data were extracted based upon the specific dates and locations of samples (larval fish or estuary locations) using the R packages ‘tidyverse’ (Wickham, 2017), ‘raster’ (Hijmans, 2019), ‘ncdf4’ (Pierce, 2019) and ‘RedaS’ (Hatzinger *et al*., 2015). For each sample, the wind direction and speed were taken as the mean of 10 pixels (15 km x 15 km) centred over the sample latitude/longitude.

To investigate the interaction between offshore advection/upwelling favourable (NE) and retention/downwelling favourable (SE) winds, we separated the onshore winds in eastern Australia into SE and NE components resulting in two variables. The SE and NE components of wind are most relevant to the transport of larvae. To calculate the magnitude of wind in a particular direction (SE or NE winds) from the known direction and speed, we converted the wind direction to radians and applied a sine function, taking the absolute value to create an effect size for wind displacement. This can be interpreted as the net movement of air moving along a specified axis (NE or SE). If the wind was directly from 45° (for NE winds; 135° for SE), then it was a full effect (1); but if it was at a slight angle, then it was reduced (<1) according to the sine function.

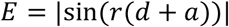

Where *E* = effect size for the wind displacement, *r* = a function to convert direction wind is going towards to radians, *d* = direction wind is going towards (°), *a* = 45° for NE calculation or 135° for SE calculation.

The effect size was then multiplied by the wind speed to get the displacement in each direction per hour and −1 to account for the data being direction towards and ensure offshore winds were negative.

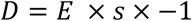

Where *D* = displacement (km h^−1^) and *s* = wind speed (km h^−1^).

Displacement values were then summed to generate a total net displacement (km) over the time span of interest. A positive net displacement means that a greater amount of air moved onto the coast than off the coast during the time period. The use of net displacement rather than other methods aligns with the original work of Dannevig (1907) and Suthers *et al*. (2020) while presenting a similar metric to Schlaefer *et al*. (2018). Henceforth, references to upwelling favourable winds (NE winds) or downwelling favourable winds (SE winds) refer to net displacement of air in either upwelling or downwelling favourable directions. While we are calculating wind displacement, rather than direct upwelling or downwelling (Ekman transport), wind is a key driver of upwelling in this region (Schaeffer *et al*., 2013).

#### Larval Fish Data

We subset the larval fish abundance data to include samples taken on the continental shelf (≤ 1000 m bathymetry; Figure 1) between 30–36° S for the period 1990 to 2016 (to match wind data availability). In order to investigate recruitment mechanisms affecting all coastally spawned fish, within each larval fish sample we focused only on taxa known to spawn coastally in this region (Table S2; Neira *et al*., 1998, Miskiewicz Unpublished Data). While this may miss some species-specific effects, we are assuming that coastally spawned taxa will respond in similar ways prior to their swimming ability improving with larval development, which facilitates movement from coastal waters into estuarine nursery habitats. To avoid any one taxon dominating the abundance of coastal larvae, the abundance of each taxon was normalised (i.e. the abundance of each family summed to 1) by transposing the dataset and using the ‘*normalize.rows()*’ function from the ‘vegetarian’ R package (Charney and Record, 2012). This created a relative abundance measure and ensured that each taxon had equal weighting during subsequent analysis, removing species specific effects. The normalised abundance of each taxon was summed together to create a single metric for analysis. Following normalisation, the normalised relative coastal abundance across all samples had a mean of 0.07 (SD = 0.11), with a maximum of 1.48 and the mean abundance was stable between sampling projects (Figure S1). By using the relative abundance of a suite of species, the possibility exists that some individual species-level effects may be missed, but during the larval stage (and without size information) we are assuming that swimming ability is poor and that all species will respond similarly to coastal winds. The larval fish sampling in Smith *et al*. (2018) was spatially inconsistent and in this region larval abundance is also generally higher in shallower areas (Hinchliffe *et al*., 2021). Therefore, to control potential bias, we calculated the distance (km) to the nearest point of the Australian mainland for each sample to use as a covariate during modelling.

**Figure 1.**
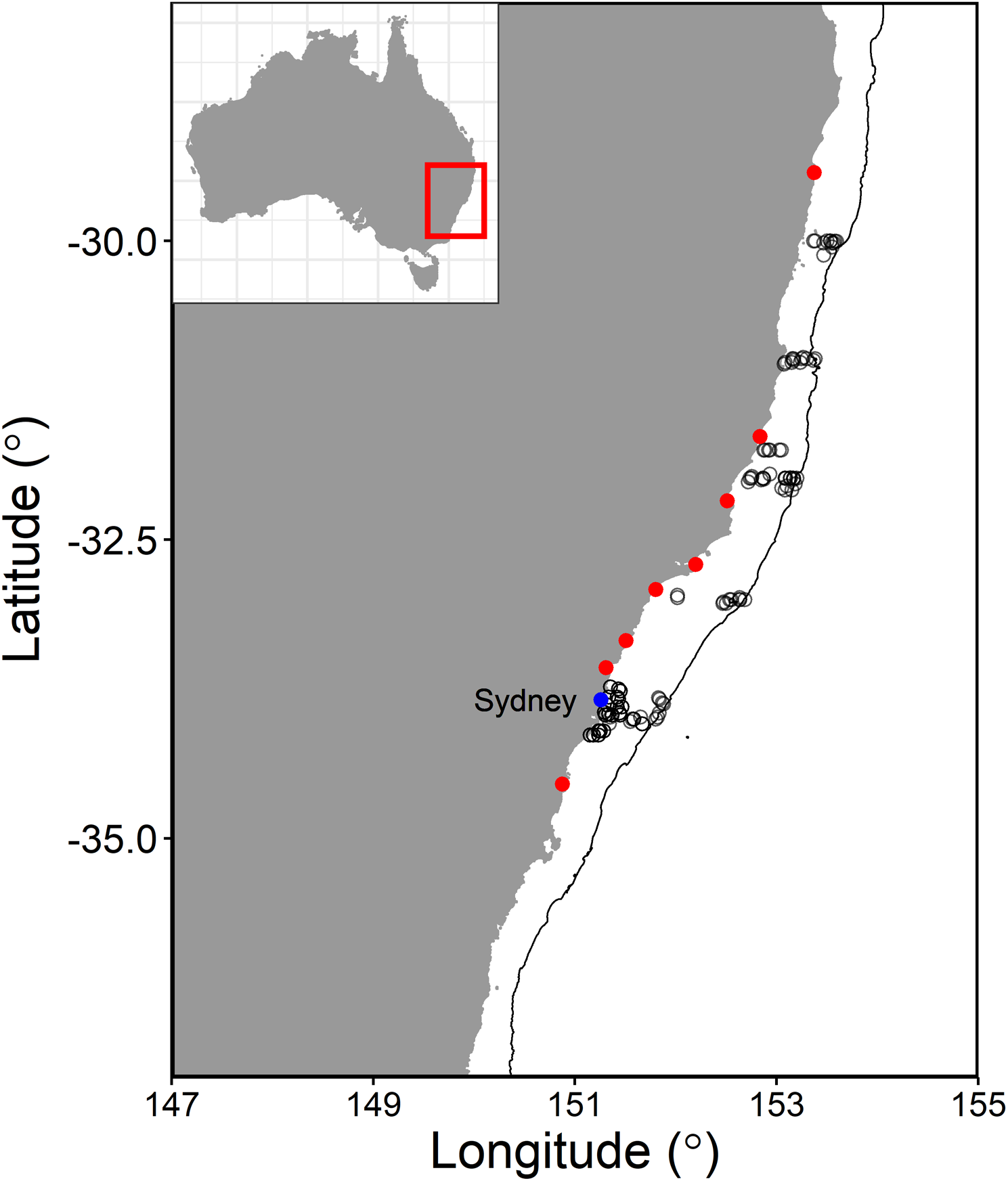
Location of southeast Australia showing the position of the larval fish samples included in the study (empty grey circles), the estuaries used in the catch-per-unit-effort analysis (filled red circles) and Sydney (filled blue circle). The black line represents the 1000 m isobath.

To separate the effects of advection from upwelling favourable (NE) winds and onshore transport from downwelling favourable (SE) winds on larval fish abundance, the net air displacement for upwelling favourable and downwelling favourable winds for each larval fish sample was calculated by summing the hourly displacement measurements of the previous 14 days. This time period was selected to quantify both potential upwelling and retention against the coast as there are often lagged effects on larval abundance through physical transport and nutrient enrichment (Dalley *et al*., 2002; Ings *et al*., 2008). While larval transport can happen over short periods of time, 14 days is the approximate period it takes for upwelling to generate secondary production (increased phytoplankton and zooplankton) in this region and therefore provide a potentially favourable environment for fish larvae to develop (Baird *et al*., 2006).

#### Commercial Catch

For our investigation, we aggregated all monthly data to an annual scale (July – June). Over the ten-year period, there were no major regulation or fisher behaviour changes within this fishery and both total catch and CPUE fluctuated despite relatively stable fishing effort within each estuary (Figures S2 – S4). We therefore proceeded under the assumption that fluctuations in annual CPUE may be a reliable proxy for fish abundance (Gillson *et al*., 2009).

To investigate the effects of upwelling and downwelling favourable winds during the spawning periods of the commercially important fish species and the effects on lagged annual CPUE, onshore winds were calculated for each estuary (located 0.15°E of the estuary mouth; Figure 1), centred, and scaled according to the above method with upwelling (NE) and downwelling (SE) favourable components. For all estuaries, there were years of relatively high and low winds (Figures S5 & S6). The net air displacement during the spawning period was determined by identifying the spawning periods of each species from published literature (Table 1). Rather than exploring multiple lags, we identified lags *a priori* by using the modal age of these species caught by gillnets in these estuaries (Gray *et al*., 2015). The modal age is the age at which the species are fully recruited to the fishery, removing any effect of gear selectivity. This age was then used to lag the spawning period winds to correspond to the most abundant (modal) age class, which are therefore most likely to show an effect of the onshore winds if they were influencing larval recruitment. As drought has previously been shown to be an important driver for this CPUE dataset (Gillson *et al*., 2009), we included drought as the number of months each estuary was ‘drought declared’ during each 12-month CPUE period based on the New South Wales Department of Primary Industries drought situation maps. Our dataset also contained three types of estuaries (barrier river, drowned river valley & barrier lagoon) which might have differing responses to drought due to their different flushing schemes (Roy *et al*., 2001).

### Statistical Analysis

#### Effects of Wind on Larval Fish Abundance

The larval fish data covered a period of 27 years (1990 – 2016) and included 1,489 larval fish samples that were used to calculate the normalised coastal abundance metric for each sample described above. A total of 60 larval fish samples (4%) contained no (zero) coastally spawned larvae. Rather than remove these samples we used a two-stage gamma hurdle model to test the effects of upwelling favourable (NE) and downwelling favourable (SE) winds on larval fish abundance. This gamma hurdle model first analyses all the data in a presence-absence method using a Bayesian binomial model with a logit link. This is followed by a Bayesian generalised linear mixed model with a gamma error distribution using a log link for the presence only data because the abundance data had a continuous, positively skewed, distribution. The model included the fixed effects of downwelling favourable winds, upwelling favourable winds and distance from the coast (km) as well as interaction terms and separate quadratic terms for both downwelling and upwelling favourable winds with interactions with distance-to-coast. Both linear and quadratic terms for the downwelling and upwelling favourable winds were included in the models because it was hypothesised that the winds would have a disproportionate effect on larval fish abundance at low or high speeds, hence a non-linear fit may be appropriate. Distance-to-coast was included as a covariate to control for the natural increase in larval abundances closer to the Australian mainland (Hinchliffe *et al*., 2021). Distance-to-coast, upwelling favourable (NE) and downwelling favourable (SE) winds were centred and standardised to assist the model fitting process and increase interpretability of the model coefficients (Schielzeth, 2010). This means a wind of 0 is interpreted as moderate (i.e. the mean) while positive values are stronger winds and negative values are weaker than the mean.

Since the Australian IMOS Larval Fish database includes data from a variety of projects which used slightly different sampling methodologies (Smith *et al*., 2018), the model also included a random intercept effect for Project. As each Project occurred in a discrete time period, this random effect also controls for any temporal inconsistencies such as a traditional year effect. The use of normalised larval abundances also creates a more consistent dataset despite the differences in sampling over time. The fitted model thus had the form:

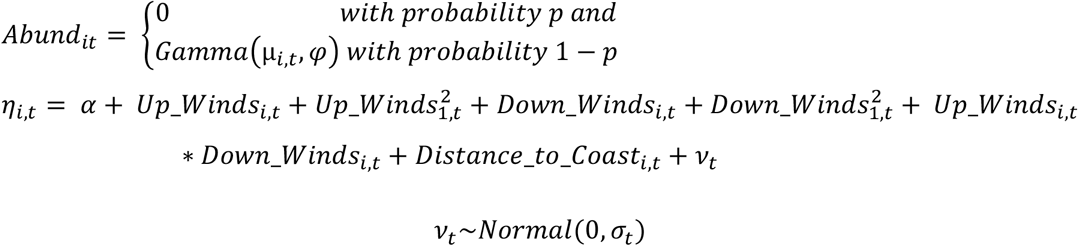

Where the probability that the relative abundance of coastally spawned larvae (*Abund*) is 0 was modelled as logit(*p*); *α* is a constant intercept; Variables relate wind and distance-to-coast variables to the abundance of coastally spawned larvae in sample *i* from project *t*. * represents an interaction term in the model. *Abund* was modelled as Gamma distributed with mean μ_*i,t*_ fitted via a log link with linear predictor *η*_*i,t*_, and shape parameter *φ*. *ν*_*t*_ is a random intercept by project with mean 0 and variance 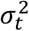.

Model parameters were estimated by MCMC using Stan (Carpenter *et al*., 2017) using the function brm() of R package ‘brms’ (Bürkner, 2018). Four parallel MCMC chains were run for 10,000 iterations (5,000 warm up and 5,000 samples each chain) and all parameter estimates were presented with their 95% Bayesian credible intervals (CI). Convergence was assessed by visually examining MCMC trace plots and assessment of the Gelman–Rubin statistic (Brooks and Gelman, 1998). The overall model fit was assessed by extracting residuals from the model and generating standard Pearson’s residual vs fitted plots and normal quantile plots, following the model checking procedure of Gillson *et al*. (2020). The model was considered stable as the chains were well mixed (Figure S7) the Gelman–Rubin test statistic < 1.01 for all parameters. The shape and hurdle parameter estimates did not overlap zero (shape 95% CI: 0.54 – 0.62, hu 95% CI: 0.13 – 0.17), thereby justifying the added model complexity in this case. There was some deviance from model fit (Figure S8), particularly in the tails of the dataset but due to the large number of samples and appropriateness of the gamma hurdle structure we proceeded with the analysis.

Default weakly informative priors were used for all parameters: improper flat priors over the reals for covariate effects, Student-*t*(*μ* = −3.1, *σ* = 2.5, *ν* = 3) for the intercept, Student-*t*(*μ* = 0, *σ* = 2.5, *ν* = 3) for the standard deviation terms, Gamma(0.01, 0.01) for *φ* and logistic(0, 1) for the zero *Abund* probability parameter. To visualise the effects of the model predictors, marginal effects were calculated using the ‘*ggeffects()*’ function in the ‘ggeffects’ R package (Lüdecke, 2018).

#### Effect of wind on estuarine fisheries catch rates

A Bayesian linear mixed model with gaussian error distribution was used to assess the effects of coastal winds on annual CPUE. The model included fixed effects of downwelling favourable winds, upwelling favourable winds (NE), drought months and estuary type, interactions between drought months and estuary type, upwelling and downwelling favourable winds as well as quadratic terms for both upwelling and downwelling favourable winds. To investigate the overall effect of onshore winds on annual CPUE and incorporate the dependency structure among observations from the same estuary or species, we used estuary as a random intercept and species as a random slope effect as part of a Bayesian linear mixed model. This controls for our data coming from the same eight estuaries over a ten-year period with species crossed with estuary. The random intercept lets each estuary have different overall level of production while the random slope effect lets each species respond differently to the environmental variables. Using the annual CPUE as the response variable, the linear mixed model was fit and assessed using the same method described above for the larval fish analysis. The fitted model thus had the form:

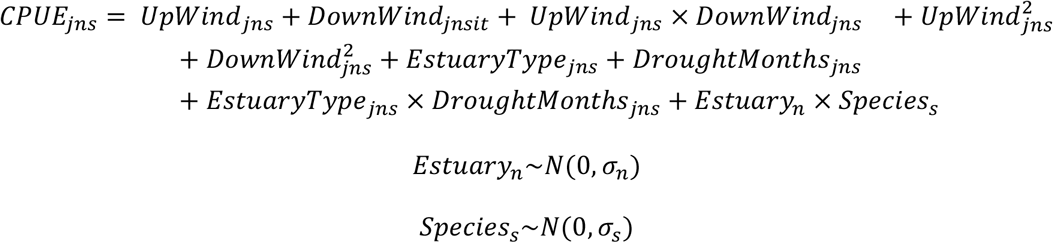

Where *CPUE*_*jns*_ is the *j*th CPUE observation in *Estuary n* for *Species s*. *Estuary*_*n*_ is the random intercept and *Species*_*s*_ is the random slope effect. *UpWind* is the standardised upwelling favourable winds, *DownWind* is the standardised downwelling favourable winds, *EstuaryType* is the type of estuary an observation was from, *DroughtMonths* is the number of months of ‘declared drought’ during the year. Uninformative flat priors were used for all variables except the overall intercept and random intercepts for *Estuary* and *Species*. The overall intercept used a Student-*t* (*μ* = 30.8, *σ* = 40.3, *ν* = 3) and the random intercepts for *Estuary* and *Species* both used Student-*t* (*μ* = 0, *σ* = 40.3, *ν* = 3). The CPUE model showed good convergence of chains with Gelman–Rubin test statistic < 1.01 for all parameters but there was some divergence in model fit towards the tails (Figures S9 & S10).

#### Historical Wind Changes

To investigate whether or not upwelling or downwelling favourable winds have changed over time in southeast Australia, we used the 20^th^ Century Reanalysis V2c dataset (1850–2014; Compo *et al*., 2015) and calculated upwelling favourable (NE) and downwelling favourable (SE) winds as described above, except net annual displacement was derived by summing all winds during a Gregorian calendar year (centred on Sydney 33.839° S 151.309° E; Figure 1). To ensure consistency with the high-resolution BARRA model used for the prior analyses, we initially tested the correlation between the BARRA model and 20^th^ Century Reanalysis V2c model using both annual displacement for upwelling favourable (NE) and downwelling favourable (SE) winds for the overlapping years (1990– 2014) centred over Sydney. There was a moderate to high correlation for both upwelling favourable (*r* = 0.599, *t*_23_ = 3.589, *P* = 0.002) and downwelling favourable (*r* = 0.697, *t*_23_ = 4.658, *P* < 0.001) winds. We then applied two separate Bayesian linear models with gaussian error distributions for upwelling favourable (NE) and downwelling favourable (SE) winds with year as a fixed effect (flat uninformative prior). Initial exploration revealed temporal autocorrelation in the first and second year of the wind time-series data as measured with the ‘*acf*’ function from the ‘stats’ R package (R Core Team, 2020). Therefore, wind data from every third year were used in the final analysis to remove the presence of temporal autocorrelation. Bayesian linear models were fit and assessed using the same method described above for the larval fish analysis, with good mixing of chains and model fit (Figures S11 – S14).

#### Interpretation and Sensitivity Analysis

For interpretation of all model outputs in this study we used the median posterior estimate for parameters and the 95% Bayesian CI. If the 95% CI did not overlap zero, we deemed a parameter important (van der Linden and Chryst, 2017). Conditional (full model) and marginal (fixed factors only) *R*^2^ values were calculated using the method of Gelman *et al*. (2019) and implemented in the function ‘*r2_bayes()’* of R package ‘performance’ (Lüdecke *et al*., 2020).

For both the larval and CPUE models, sensitivity analyses were performed on the lag times used to assess the robustness of our findings. This involved running the models multiple times while altering the duration of the lag period. For the larval models, we varied the lead up time for winds by 9 – 20 days prior to sampling and for the CPUE models we simulated altering the lag by ± 2 years from the identified modal age. This is important for the CPUE models as there are potential influences in the data due to the harvesting of multiple age classes. As we chose the modal age from gillnet sampling, our original lag should have the strongest effect, and if multiple age classes are present in a sample, then the effects of wind should be similar but potentially weaker if an age class is less abundant.

## Results

### Coastally Spawned Larval Fish

A total of 175,112 larval fish from 132 coastal spawning taxa were present in 1,489 observations on the continental shelf between 30 and 36° S. Examining the winds 14 days prior to sampling revealed evidence of an interaction between distance-to-the-coast, upwelling favourable (NE) wind and downwelling favourable winds (Estimate = −0.48, 95% CI: −0.85 – −0.13; Figure 2). This interaction can be interpreted as upwelling and downwelling favourable winds interact with each other in different ways depending on the distance a sample is taken from the coast. An overall decline in abundance was observed with increasing distance from the coast (Figure 3). Upwelling favourable winds had a nonlinear effect on abundance where when upwelling favourable winds increased from below average to moderate amounts (0 on standardised axis), larval abundance increased but then as the winds strengthened further, abundance decreased (Figure 3). This resulted in an optimum threshold where larvae were most abundant following moderate amounts of upwelling favourable wind. Downwelling favourable winds showed a linear positive effect on abundance (Figure 3). Both the upwelling and downwelling favourable wind relationships were strongest near the coast and the relationships became less defined with increasing distance from the coast (Figure S15).

**Figure 2.**
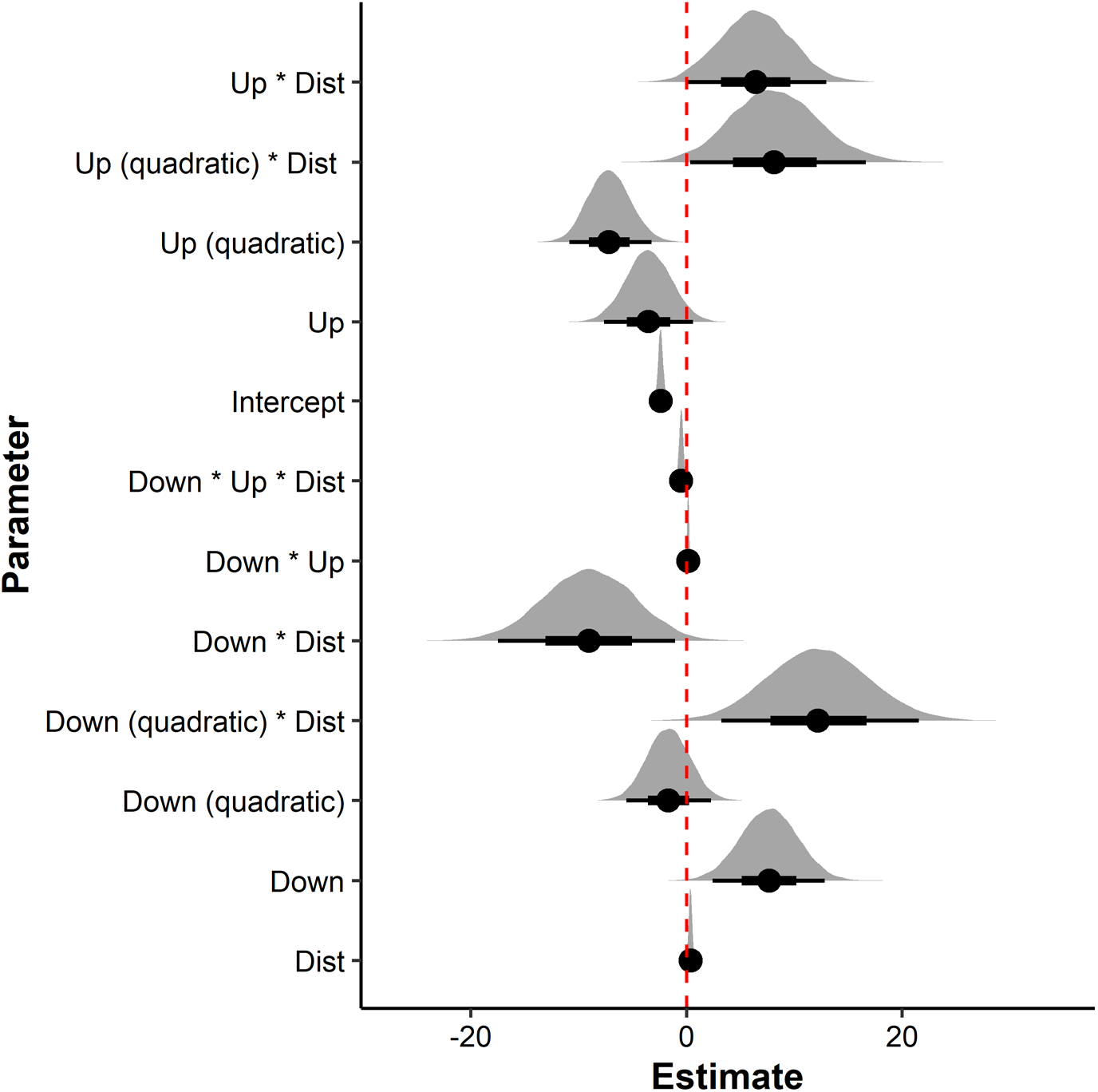
Caterpillar plots showing the Bayesian parameter estimates for the coastally spawned larval fish abundance models based upon winds in the 14 days prior to sampling. Points show the estimate of the parameter with the horizontal bar representing the 66% (thick bar) and 95% (thin bar) credible interval of the estimate. Model terms include distance-to-coast (Dist), upwelling favourable winds (Up) and downwelling favourable winds (Down). Parameter estimates are deemed important if the 95% credible interval does not cross the dashed red line which marks an estimate of zero. For exact estimate values see Table S3.

**Figure 3.**
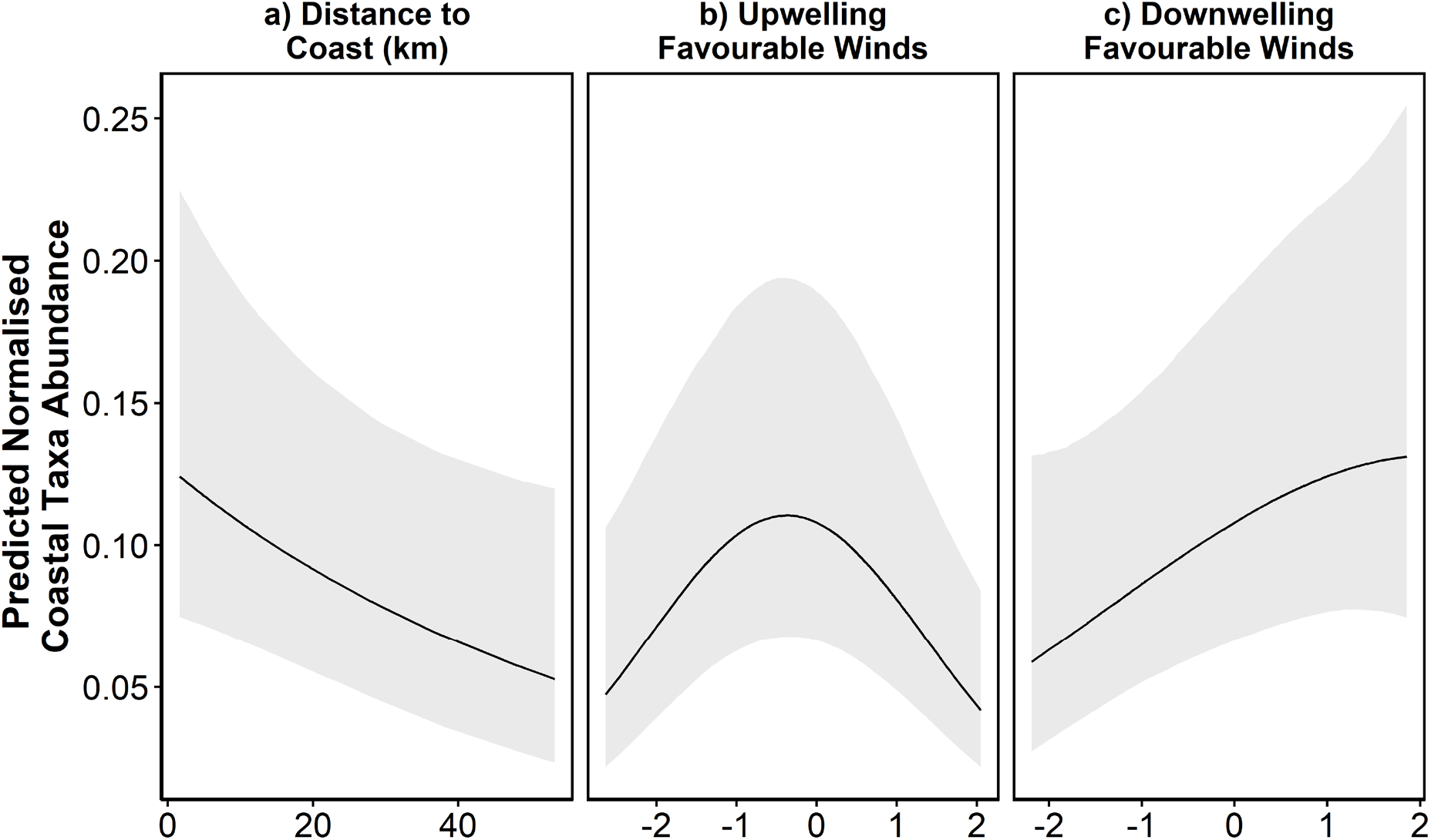
Visualisation of the predicted normalised coastal larval abundance based on the generalised linear mixed model for winds 14 days prior to sampling: a) the effect of distance from the coast, based upon mean (0 on the standardised scale) upwelling and downwelling favourable winds with shading showing the 95% credible interval; b) the effect of upwelling favourable winds; and c) the effect of downwelling favourable winds. Predictions for each variable were made while holding all other variables within the model to mean values (0 for wind variables and 10km for distance from the coast). For the winds, 0 represents mean winds, with 1 and −1 representing 1 and −1 SD from the mean, respectively.

Our 14-day wind model explained a small amount of variance in larval fish abundance (conditional *R*^2^ = 0.059, marginal R^*2*^ = 0.044), and the sensitivity analysis of the wind lead up times in the larval fish model showed that estimates of most effect sizes did not clearly change between 9 – 21 days lead up (overlapping error bars; Figure S16). Two changes were evident in the sensitivity analysis: 1) the interactions between upwelling favourable winds and downwelling favourable winds became inconsequential when a 9 – 12 day lag was tested (error bars overlaps 0); and 2) the effect size of interaction between upwelling favourable winds, downwelling favourable winds and distance-to-coast became smaller at a 9 day lag, although still remained important and negative.

### Commercial Estuarine Catch Rates and Historical Winds

The multi-species model performed well (conditional *R*^2^ = 0.822, marginal *R^2^* = 0.506) and showed strong evidence for a negative effect of upwelling favourable winds during the spawning period on CPUE (Estimate: −230.48, 95% CI: −374.58 – −85.05; Figures 4 & 5). There was no evidence of any effect from downwelling favourable winds on CPUE (Estimate: −88.62, 95% CI: −227.90 – 50.63; Figure 4).

**Figure 4.**
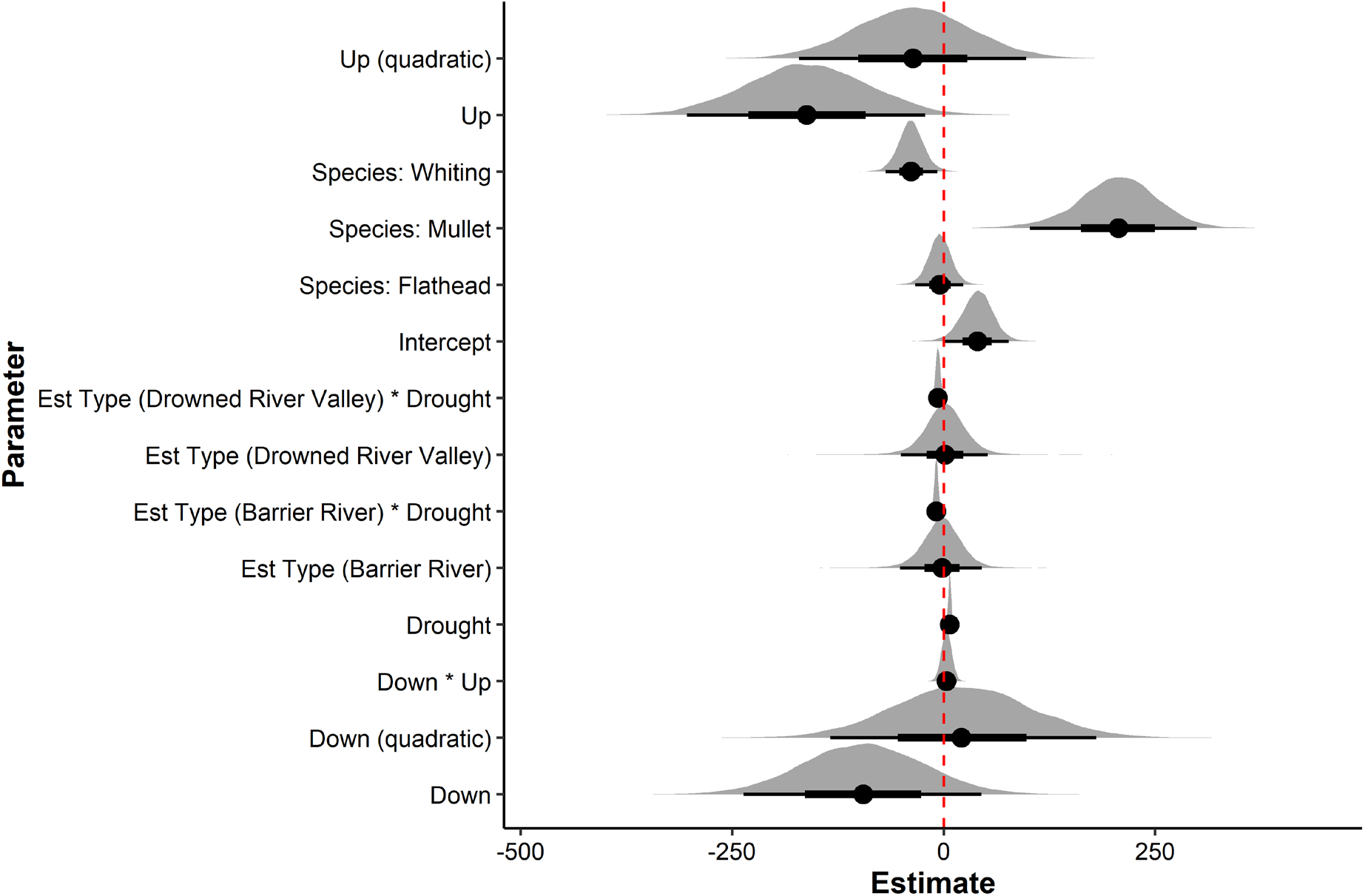
Caterpillar plot showing the Bayesian parameter estimates for the multi-species catch-per-unit-effort (CPUE) generalised linear mixed model. Model terms include upwelling favourable winds (Up), downwelling favourable winds (Down) and estuary type (Est Type). Points show the estimate of the parameter with the horizontal bar representing the 66% (thick bar) and 95% (thin bar) credible intervals (CI) of the estimate. The dashed red line marks an estimate of zero. The 95% CI does not cross zero for Up Winds, Species: Whiting, Species: Mullet, Drought Months, Estuary Type: Drowned River Valley * Drought Months and Estuary Type: Barrier River * Drought Months. For exact estimate values see Table S4.

**Figure 5.**
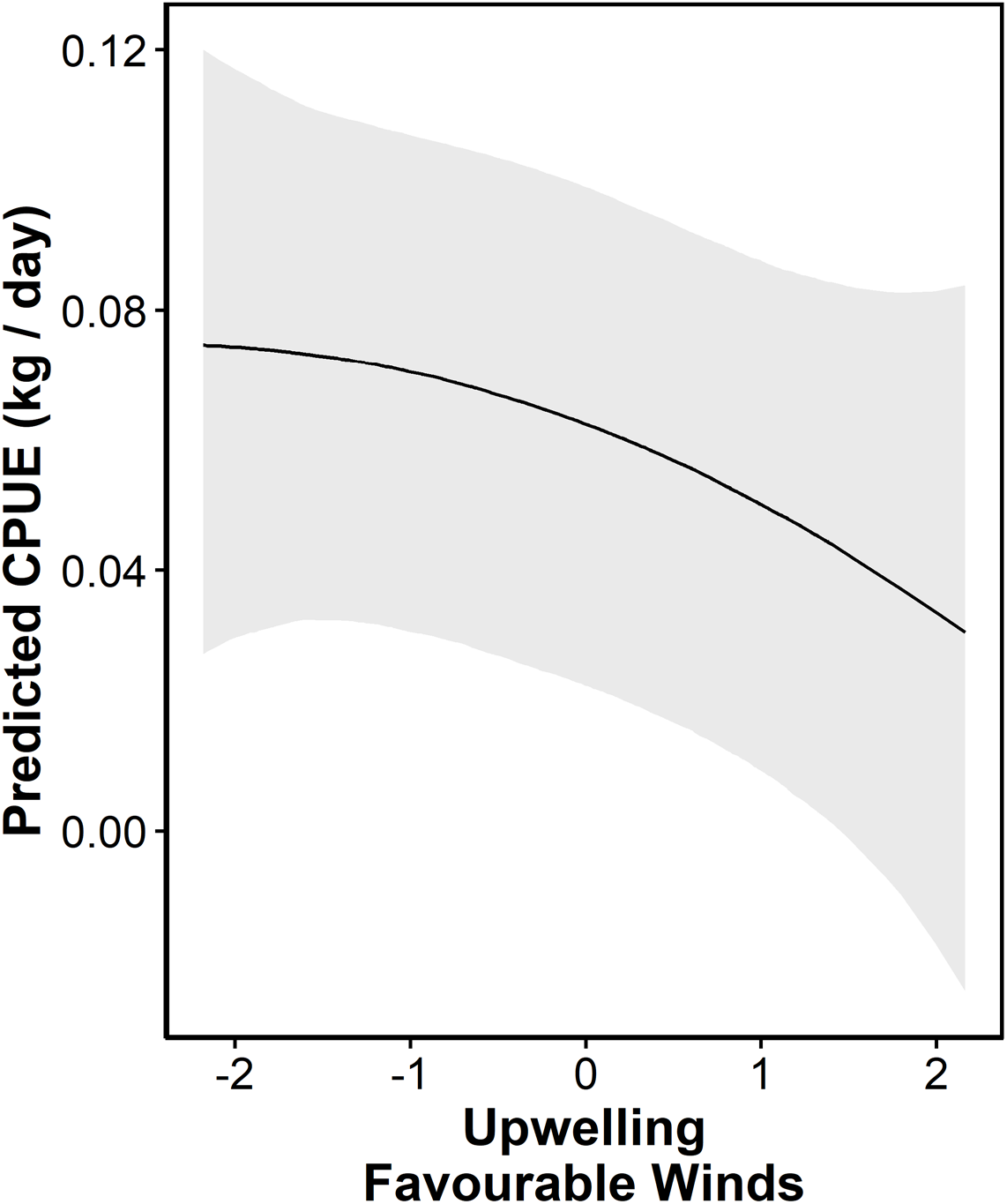
Visualisation of predicted catch-per-unit-effort (CPUE) based upon upwelling favourable winds during the spawning period as predicted by the results of the CPUE Bayesian linear mixed model. The grey area represents the 95% credible interval. There were no detectable effects of downwelling favourable winds and the effects of drought are shown in the supplementary material (Figure S17). Predictions were made while holding all other variables within the model to mean values. Note due to the random slope and intercept effects in the model, the actual y scale is relative.

Aside from the wind effects, there was evidence of a positive effect of drought on CPUE (Estimate = 8.16, 95% CI: 4.56 – 11.70) although this effect was variable depending on estuary type. Barrier Lagoon estuaries had a more positive response to drought compared with Barrier Rivers and Drowned River Valleys (Figure S17).

The sensitivity analysis of the wind lag times in the CPUE model showed that altering the lag times only influenced the wind effects with the standard error of all other estimates overlapping the lag change 0 estimates for all the lags considered (Figure S18). When the lag was shortened by one year or increased by two years, there was an interaction evident between upwelling favourable winds and downwelling favourable winds (Figure S18). The effect size of upwelling favourable winds varied with lag time. The effect became inconsequential when the lag was shortened by two years while a 1 year longer lag had a stronger effect on CPUE compared to a 1 year shorter lag. The effect of altering the lag time of downwelling favourable winds was more variable with a non-linear pattern occurring if the lag was increased by two years. If the lag was reduced by 1 year, the effect of downwelling favourable winds became inconsequential.

Between 1850 and 2014, there was clear evidence of an increase in upwelling favourable winds (Estimate of annual change: 52.86, 95% CI: 19.00 – 85.59; Figure 6) and a decline in onshore transport causing downwelling favourable winds (Estimate of annual change: −40.93, 95% CI: −78.18 – −3.71; Figure 6). The upwelling favourable winds shifted from a negative net displacement to a positive net displacement (Figure 6), meaning that along the 45° axis the prevalence of upwelling favourable winds has increased over time. The downwelling favourable winds remained net positive but declined to approximately half of their initial levels (Figure 6).

**Figure 6.**
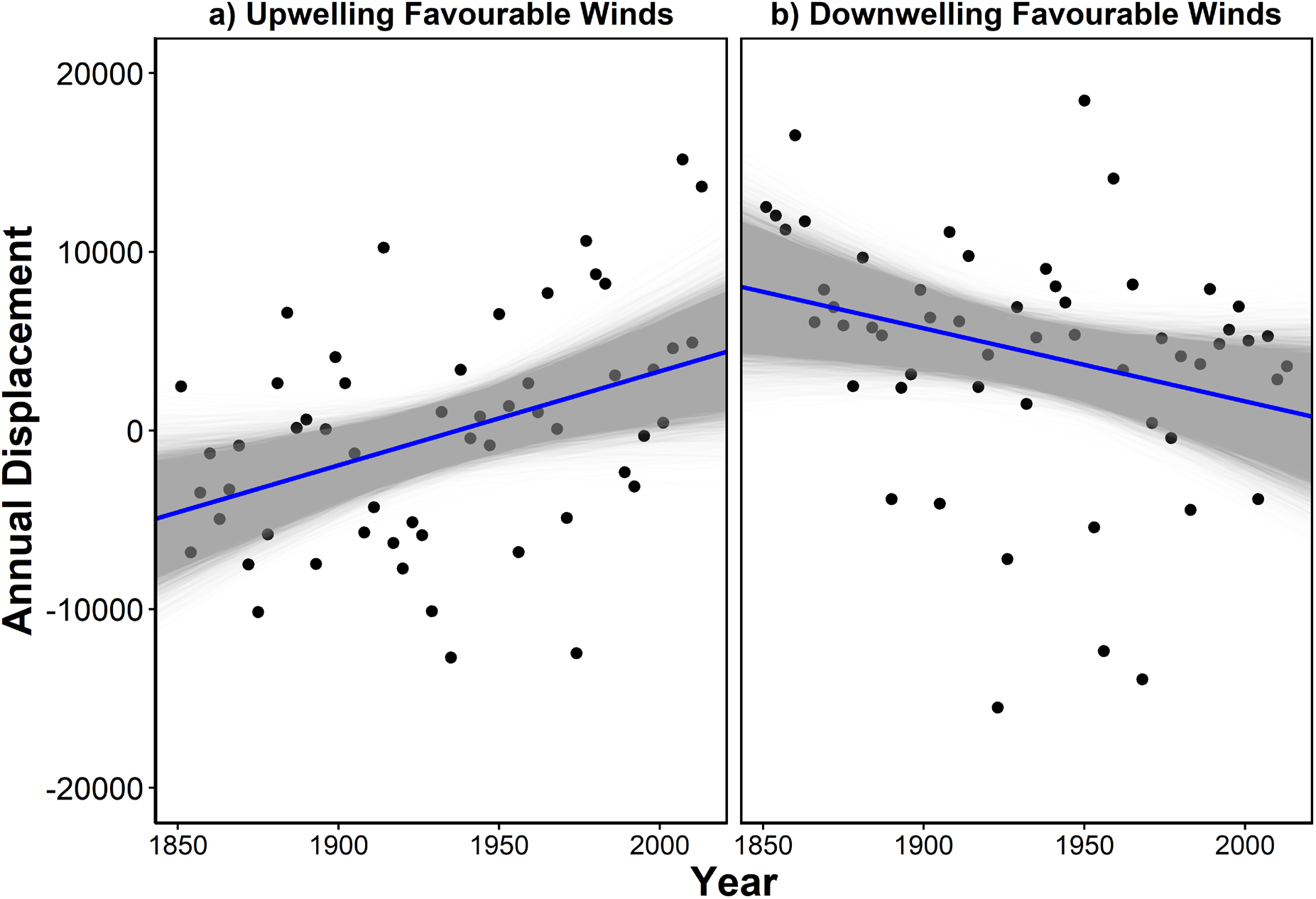
Temporal change in annual net displacement in upwelling (from northeast; Estimate of annual change: 52.86, 95% credible interval: 19.00 – 85.59) and downwelling favourable winds (from southeast; Estimate of annual change: −40.93, 95% credible interval: −78.18 – −3.71) between 1850 and 2014. The grey lines show 20,000 posterior sample estimates with the median trend line in blue. Only every third year of annual net displacement data was included in the models to account for temporal autocorrelation in the wind time-series data.

## Discussion

By combining three datasets from a larval fish database, a high-resolution wind re-analysis model and commercial CPUE time-series data, we have demonstrated that coastal winds may influence the near-shore abundance of coastally spawned taxa (Figures 2 & 3), potentially resulting in detectable effects on commercial fisheries catch rates (Figures 4 & 5). While we initially expected samples collected following strong onshore winds to have the highest abundance of coastal larvae, we found both upwelling and downwelling favourable winds are important for fish larvae. Moderate amounts of upwelling favourable winds potentially increased planktonic production (Armbrecht *et al*., 2014), although above average upwelling favourable winds negatively affected coastal larvae abundance, perhaps due to offshore advection. Strong downwelling favourable winds had a positive influence on the abundance of coastally spawned larvae, likely driving onshore transport which may facilitate estuarine recruitment (Agostini and Bakun, 2002). Using lagged winds from each species’ spawning season, we showed a detectable negative effect of strong upwelling favourable winds on CPUE, possibly due to the advection of larvae away from favourable habitat. This was a similar effect to that observed in the larval fish with strong upwelling favourable wind being correlated with low relative abundance of coastal larvae. We found no evidence of the hypothesised positive effect of downwelling favourable winds in the CPUE analysis. The high variance explained in the CPUE models (conditional *R^2^* = 0.82) in the present study indicates that incorporating wind from the spawning period into future recruitment models might improve forecasts of estuarine fisheries CPUE. This is important as we have demonstrated that coastal winds have changed since 1850 and are expected to continue changing in the future (Bakun *et al*., 2010; Sydeman *et al*., 2014).

### A proposed recruitment mechanism

We showed a negative effect of upwelling favourable wind on larval fish abundance when it was above average strength. This is likely caused by offshore transport as strong upwelling drives advection away from the coast and juvenile estuarine and nearshore habitat. It is also recognised that over an approximate 14 day cycle, upwelling can generate increased nutrients and chlorophyll at the surface which may in turn flow into the lower trophic levels (Gasol *et al*., 1997; Buesa, 2019). This upwelling may increase prey availability for larval fish, thereby increasing growth and survival rates (Zenitani *et al*., 2007). Onshore transport from downwelling favourable winds is important as it retains larvae near the coast. Based upon previous research, the positive effects of this retention are most visible after upwelling preconditions the ecosystem with nutrients, thereby creating a favourable environment near juvenile habitat (Rykaczewski and Checkley, 2008).

Our sensitivity analysis showed a consistently positive effect of downwelling favourable winds regardless of lag time. We believe this supports our hypothesis that onshore transport may increase recruitment into estuaries by larval fishes as they are geographically closer and stochastic dispersal will be reduced (Bruno *et al*., 2018).

### Commercial Estuarine Fisheries

Consistent with Dannevig’s (1907) observation, we found coastal winds during the spawning period can influence commercial fisheries catch rates. This is consistent with the relationships we found between coastal winds and the abundance of coastally spawned larval fish. However, likely due to the coarse temporal resolution of the CPUE data, there was no evidence of a positive effect of downwelling favourable winds which represent more immediate transport effects. Based on the demonstrated effects of these winds on coastal larval fish abundance, the events most likely to contribute to successful recruitment would be short periods of upwelling (to precondition the area and generate larval prey), followed by periods of onshore transport (downwelling). We propose our results are showing a negative effect of upwelling favourable winds on CPUE because offshore transport is increased during strong upwelling, which negatively affects recruitment. As our analysis is correlative rather than a manipulative experiment, the exact mechanisms underlying the relationship between upwelling favourable winds and CPUE cannot be pinpointed. Temperature mediated changes in growth and survival are likely to be a key driver of this relationship. Upwelling favourable winds cause upwelling which draws colder water up from depth (Schaeffer *et al*., 2013). If larvae are not advected offshore with the displaced surface water they would remain in a much cooler habitat which would likely result in slower growth, potentially increasing risk of predation (Pepin, 1991; Buckley *et al*., 2008). To understand possible temperature dynamics, future work should potentially consider incorporating water temperature into studies of both larval abundance and growth as well as commercial catch data.

The generally weak positive effect of drought on CPUE (with different effects in some types of estuaries) agrees with previous research using this dataset that found monthly bream CPUE increased during periods of drought (Gillson *et al*., 2009). Despite this, previous research also showed that the monthly CPUE of other species generally declined during periods of drought suggesting that the effect of drought on CPUE needs further investigation (Gillson *et al*., 2009). We believe the contrasting results in our study compared to Gillson *et al*. (2009) are the result of using different temporal scales, with shorter timescales likely more representative of changes in both fish and fisher behaviour (Gillson *et al*., 2009).

The sensitivity analysis conducted on the lag times between the winds and year of capture identified that the modal ages show the strongest effects of coastal winds, as would be expected with ages (represented by lags) which are less common in the harvest having smaller effect sizes. In the future, it may be possible to conduct more detailed analyses of single species by identifying bottlenecks in the lifecycle including recruitment, which may be affected by coastal winds.

### Historical changes in onshore winds

Since 1850, upwelling favourable winds have increased while downwelling favourable winds have decreased in the southeast Australian region. While the direct cause of these changes is uncertain, it is possible this has been a response to global climate change. For example, intensification of surface winds have been attributed to a decline in ozone around Antarctica, which resulted in large scale changes in southern hemisphere winds including over Australia (Cai, 2006). The demonstrated changes in wind in southeast Australia would likely have reduced the onshore transport of larval fish, impacting recruitment to estuaries. Increased upwelling and decreased downwelling would increase prey availability but reduce retention near the coast, potentially transporting larvae further away from estuaries, due to similar processes as those described in Sorte (2013). This change in onshore transport may interact with the strengthening EAC which is also pushing further poleward (Suthers *et al*., 2011; Wu *et al*., 2012). If there are differences in available juvenile habitat in either cross-shelf or along-shelf directions, then the recruitment of larvae may be impacted (Sorte, 2013).

Despite this, very few fish species spawn all year round, and it is possible that seasonal changes in wind may be more important than annual changes. The temporal analysis presented here does not test alternating upwelling and downwelling favourable winds, and it would be useful to calculate a metric that encompasses the alternating nature of upwelling and downwelling favouring winds. A possible approach could be to document the number and duration of upwelling and downwelling events.

Previous research has indicated that the intensity of wind driven upwelling is likely to increase in most regions, with the exception of southwestern Africa (Bakun *et al*., 2010). This agrees with our finding that upwelling favourable winds have historically increased in southeast Australia. While this may result in increased fisheries production by increasing nutrient availability, it is a complex system in which changing winds could have a multitude of effects including increasing productivity and nutrient concentrations of source waters, regional changes in stratification and basin-scale changes in thermocline structure (Bakun *et al*., 2010).

### Limitations of this study

While our study successfully combined three datasets to assess the effects of wind on larval fish retention and recruitment, it does have limitations. We used correlative analyses and several potential explanatory variables were not included in our models, which may have captured additional variance in either larval fish abundance or commercial fisheries catch rates. These include varying population spawning biomasses, water temperature, oceanographic currents, improved larval swimming ability with development or the effects of density dependence, all of which influence spawning and/or recruitment (Ottersen and Sundby, 1995; Schilling *et al*., 2020).

The metric of wind calculated in the present study (adjusted wind speed along certain vectors) is different to conventional analyses which typically use speed and direction vectors and directly calculate movement along an axis (e.g. Schlaefer et al., 2018). Our method applies a non-linear transformation based upon the shape of a sine wave from −1 to 1, which results in a weaker adjustment on directions closer to the direction of interest and a harsher adjustment on more perpendicular directions. We believe this transformation is more appropriate where the coastline is not uniform but has minor deviations in orientation, relevant to the dispersal of larvae.

An important limitation of the present study was that by using a historical larval dataset, which was collected opportunistically, we could not directly quantify the movement or survival of larval fish during wind events and instead we could only look at abundances at the time of sampling while controlling for variation such as distance from coast and the individual project which collected the data. A more robust analysis would be to structure a survey that repeatedly samples in a grid pattern during wind events and directly quantifies the movement of larvae. As no larval fish size data were available, swimming ability was not considered, which is known to be an important factor influencing larval movement (Leis, 2007; Drake *et al*., 2018). While this could potentially bias analyses, we considered the swimming ability of larvae within each sample to be representative of a normal distribution (modelled as part of the random project effect) and any strong swimmers would have likely avoided capture by the slow plankton net.

The CPUE analysis has several key limitations which may obscure the effects of wind during the larval period. The most important of these is the duration between recruitment and commercial harvest, which can be up to 7 years in our study. This is a large time interval during which there may be many sources of mortality or population limitation. A potential limiting factor in estuaries is habitat availability, if there is insufficient suitable habitat to support recruiting larvae as juveniles, then the total population of the species will be limited by habitat rather than larval recruitment. While no information is available on potential habitat limitation in the estuaries we studied, in nearby estuaries closed to commercial fishing there is some evidence of habitat limitation (Folpp *et al*., 2020). The other limitation is that multiple age classes are harvested at a time and by choosing a modal age (based upon 2008-2010 harvest; Gray *et al*. 2015), any effects with the influence of other ages are blurred. While not possible for this study, a more rigorous approach would have been to only use the CPUE of the modal age class being harvested rather than the total CPUE. This blurring effect may be a contributor to the variation observed in the CPUE sensitivity analysis (Figure S18). The present study also did not investigate the abundance of larval fish entering estuaries (Boehlert and Mundy, 1988), which would be an important metric to confirm increased recruitment following favourable wind conditions. By expanding investigations beyond the larval period, fish recruitment mechanisms in this region might be better understood.

### Conclusions

Various studies have demonstrated positive effects (Nelson *et al*., 1977) and negative effects (Parrish *et al*., 1981; Takeshige *et al*., 2013) of wind-driven Ekman transport of larval fish for estuarine recruitment. Despite the low variance explained in the larval fish analysis, potentially caused by the large database containing samples taken over large temporal and spatial scales, the present study is the first to show a correlation between coastal winds and fish larvae, and detect a similar corresponding effect on the commercial fishery. We suggest a recruitment mechanism involving wind-driven coastal enrichment and larval retention during the spawning season, which is evident in commercial fisheries catch rates. This mechanism involves increased productivity from wind-driven upwelling and increased retention near the coastline, which work in tandem to increase overall recruitment of larvae into estuaries. These recruitment dynamics were potentially detected in commercial estuarine fisheries data when appropriate lags are applied and by incorporating spawning period winds into recruitment models, may improve predictions of commercial fisheries catch rates. As climate change is altering wind patterns, it is likely that wind driven ocean dynamics will continue to vary and it is recommended that scientists and management consider potential changes in recruitment.

## Acknowledgements

This study was prompted by Dennis Reid (Australian Museum) who brought to our attention Harald Dannevig’s original insight on the effect of coastal winds on estuarine fisheries. Support for the 20^th^ Century Reanalysis Project version 2c dataset is provided by the U.S. Department of Energy, Office of Science Biological and Environmental Research (BER), and by the National Oceanic and Atmospheric Administration Climate Program Office. HTS was supported by the UNSW Network Laboratory for Ocean Collaboration and NSW Research, Attraction and Acceleration Program. CH is supported by an Australian Government Research Training Program Scholarship. Larval fish data were sourced from the Integrated Marine Observing System (IMOS), an initiative of the Australian Government being conducted as part of the National Collaborative Research Infrastructure Strategy and the Super Science Initiative. This research includes computations using the computational cluster Katana supported by Research Technology Services at UNSW Sydney. We thank the anonymous reviewers for their constructive comments that improved this manuscript.

## Author contributions

HTS, IMS, AM & JPG collected the data, IMS conceived the ideas, HTS, CH, JPG & AM designed the analysis, HTS performed the analysis and led the writing of the manuscript. All authors contributed critically to the drafts and gave final approval for publication.

## Data Availability Statement

The larval fish data is freely available from the Australian Ocean Data Network https://portal.aodn.org.au/ (Smith *et al*., 2018). The BARRA wind model is freely available upon request to the Australian Bureau of Meteorology (Su *et al*., 2019). 20^th^ Century Reanalysis V2c data provided by the NOAA/OAR/ESRL PSL, Boulder, Colorado, USA, from their Web site at https://psl.noaa.gov/ (Compo *et al*., 2015). NSW Commercial Fisheries Catch Data is available upon request to the NSW Department of Primary Industries Fisheries. Extracted data and code used in this analysis is available at https://github.com/HaydenSchilling/Wind-and-Fisheries.

