## Supplementary Material for "Coastal winds and larval fish abundance indicate a recruitment mechanism for southeast Australian estuarine fisheries"

**Supplementary Material for Schilling et al 2021**

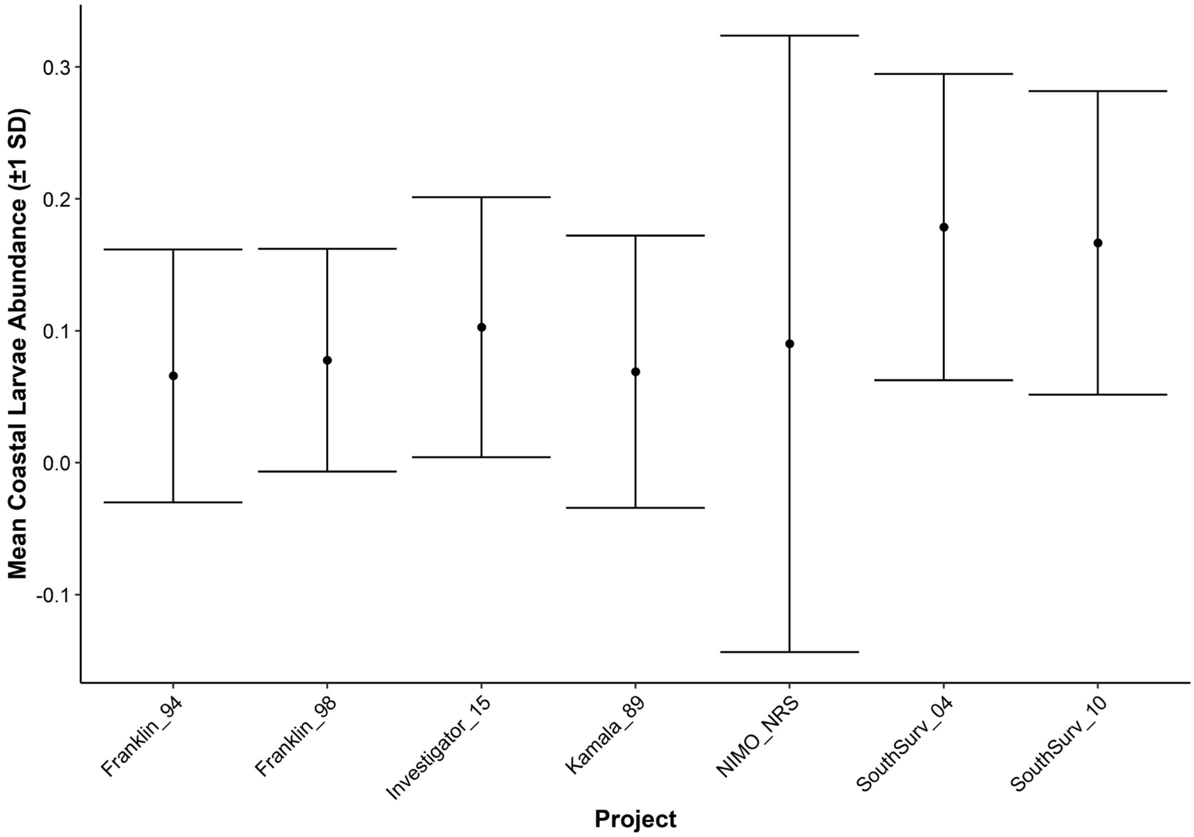


**Figure S1** Mean normalised coastal larval fish abundance among sampling projects from the Australian Integrated Marine Observing System (IMOS) Larval Fish Database (Smith *et al.*, 2018). As projects were discrete periods of time, this also shows a relative stability in coastal larval abundance through time. Projects include two *RV Franklin* voyages (1994 & 1998), One voyage on the *RV Kamala* (1989-1990), two voyages on the *RV Southern Surveyor* (2004, 2010), one voyage on the *RV Investigator* (2015) and the ongoing IMOS Larval Fish monitoring program (shown as NIMO_NRS; 2014 – 2016).

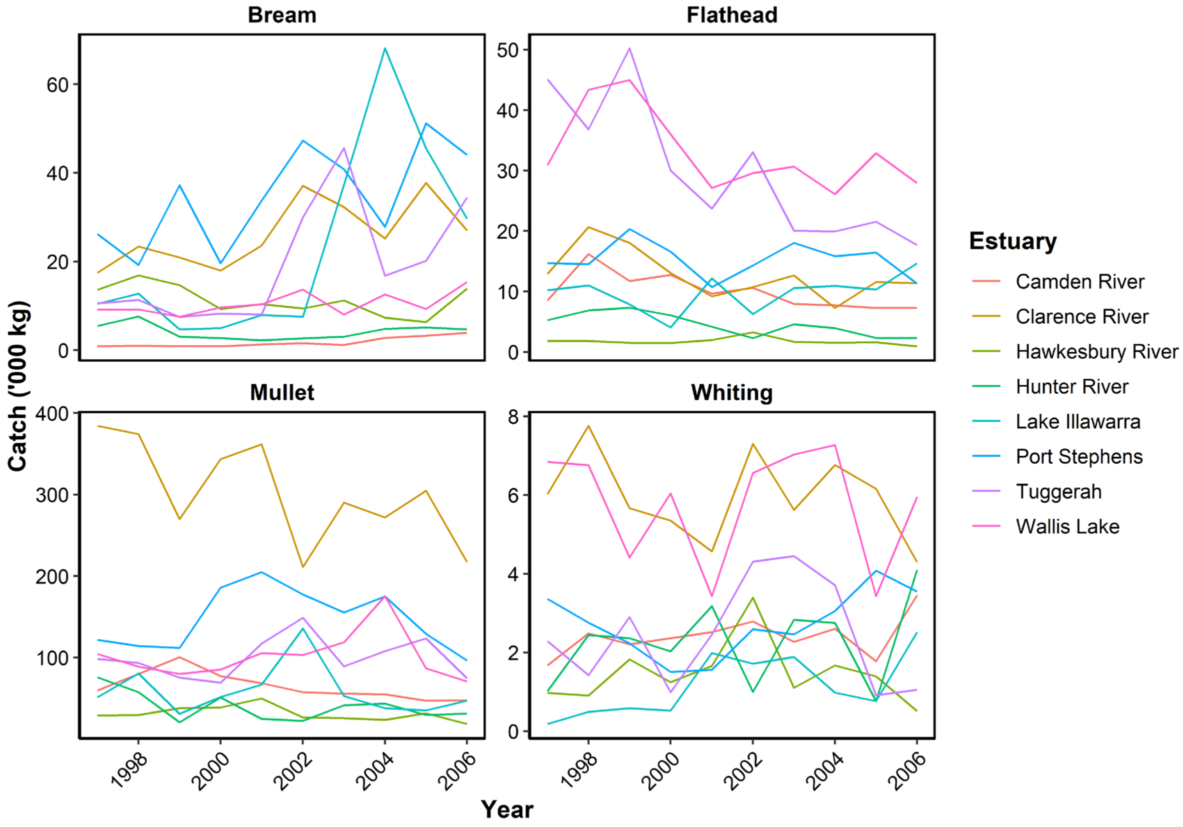


**Figure S2** Temporal trends in total annual catch ('000 kg) for four species caught in gillnet fisheries on eight estuaries from July 1997 to June 2007. Note the varying scale on the y-axis.

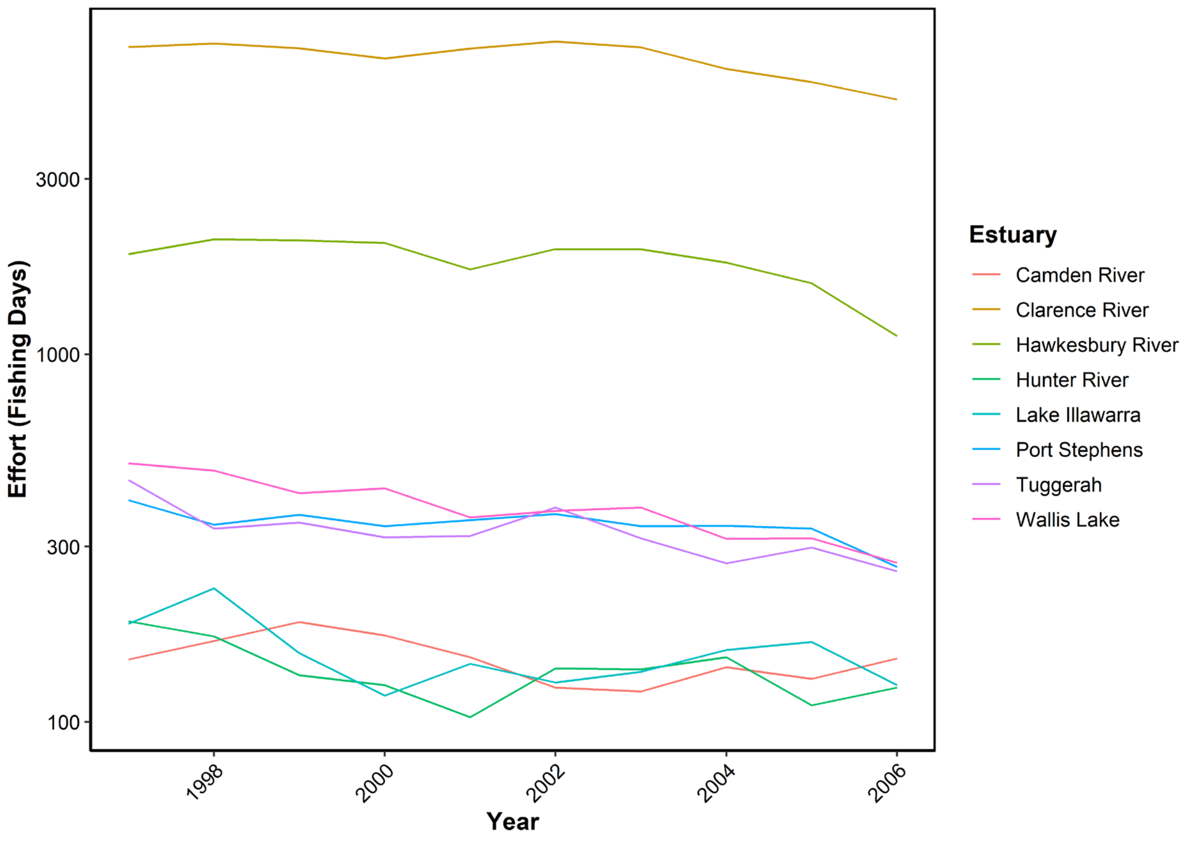


**Figure S3** Temporal trends in total annual fishing effort (days) for gillnet fisheries on eight estuaries from July 1997 to June 2007. Note the $log_{10}$ y-axis.

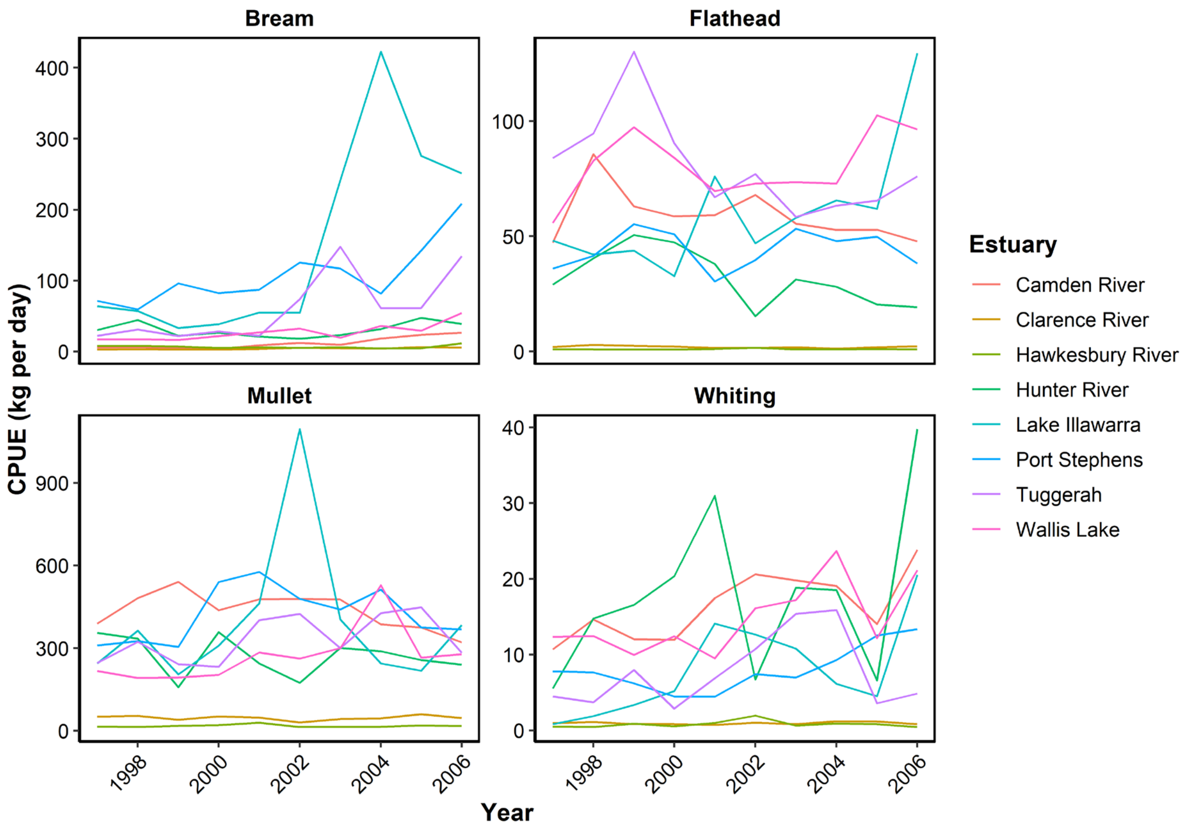


**Figure S4** Temporal trends in annual Catch-Per-Unit-Effort (CPUE; kg day$^{-1}$) for four fish species caught in gillnet fisheries on eight estuaries from July 1997 to June 2007. Note the varying scale on the y-axis.

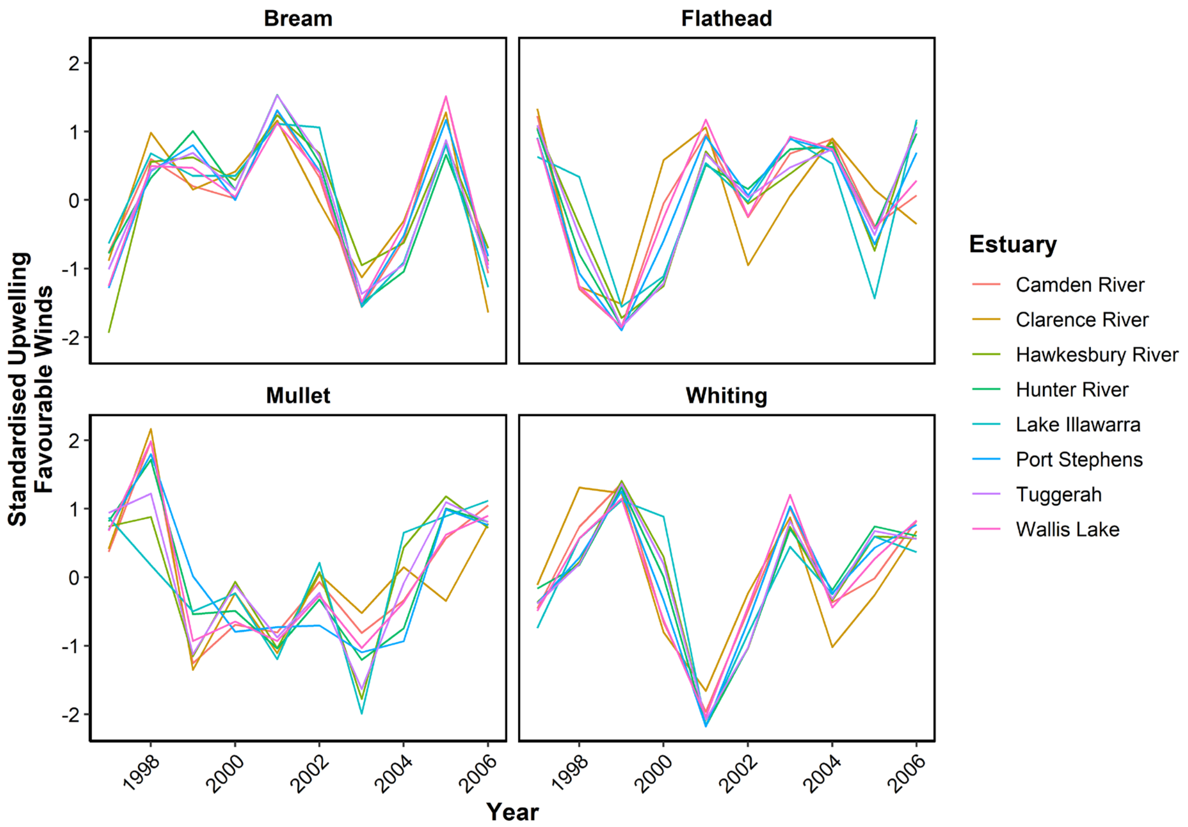


**Figure S5** Temporal trends in lagged standardised upwelling favourable winds for four fish species caught in gillnet fisheries on eight estuaries from July 1997 to June 2007. Lags are shown in Table 1.

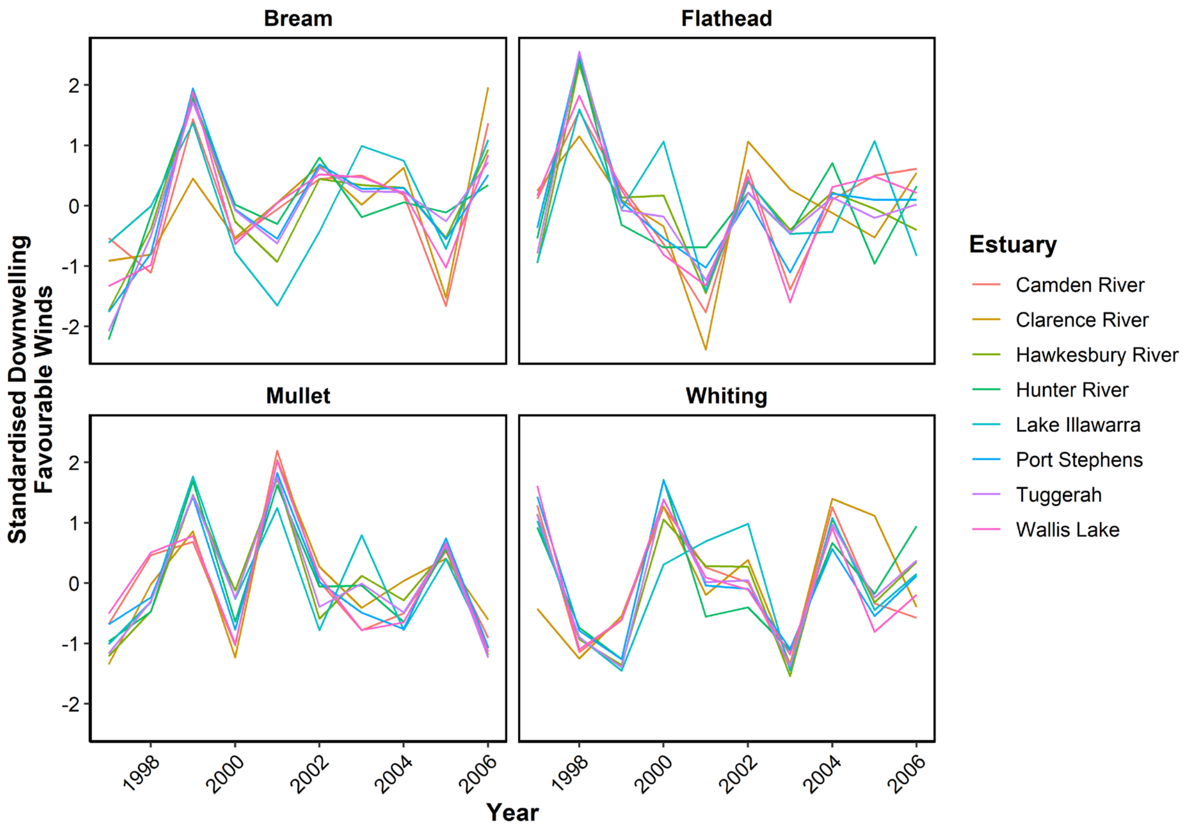


**Figure S6** Temporal trends in lagged standardised downwelling favourable winds for four fish species caught in gillnet fisheries on eight estuaries from July 1997 to June 2007. Lags are shown in Table 1.

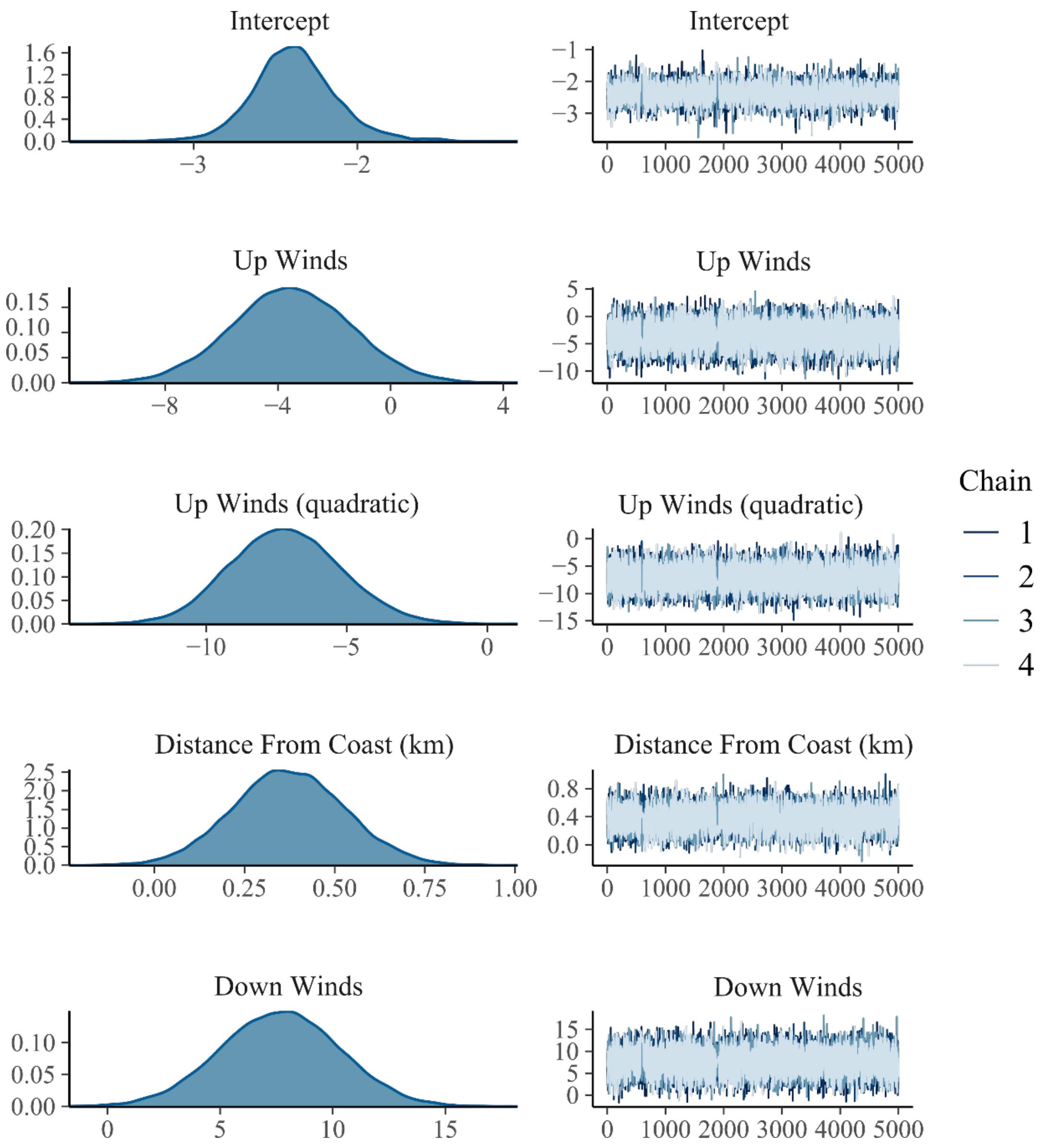


**Figure S7.1** Diagnostic plots of the parameters in the 14 day coastal larval fish Bayesian mixed model. Left hand panels show the distribution of the posterior estimates from 10,000 iterations. Right hand panels show the trace plots for the 4 chains post warm-up. Continued over the page.

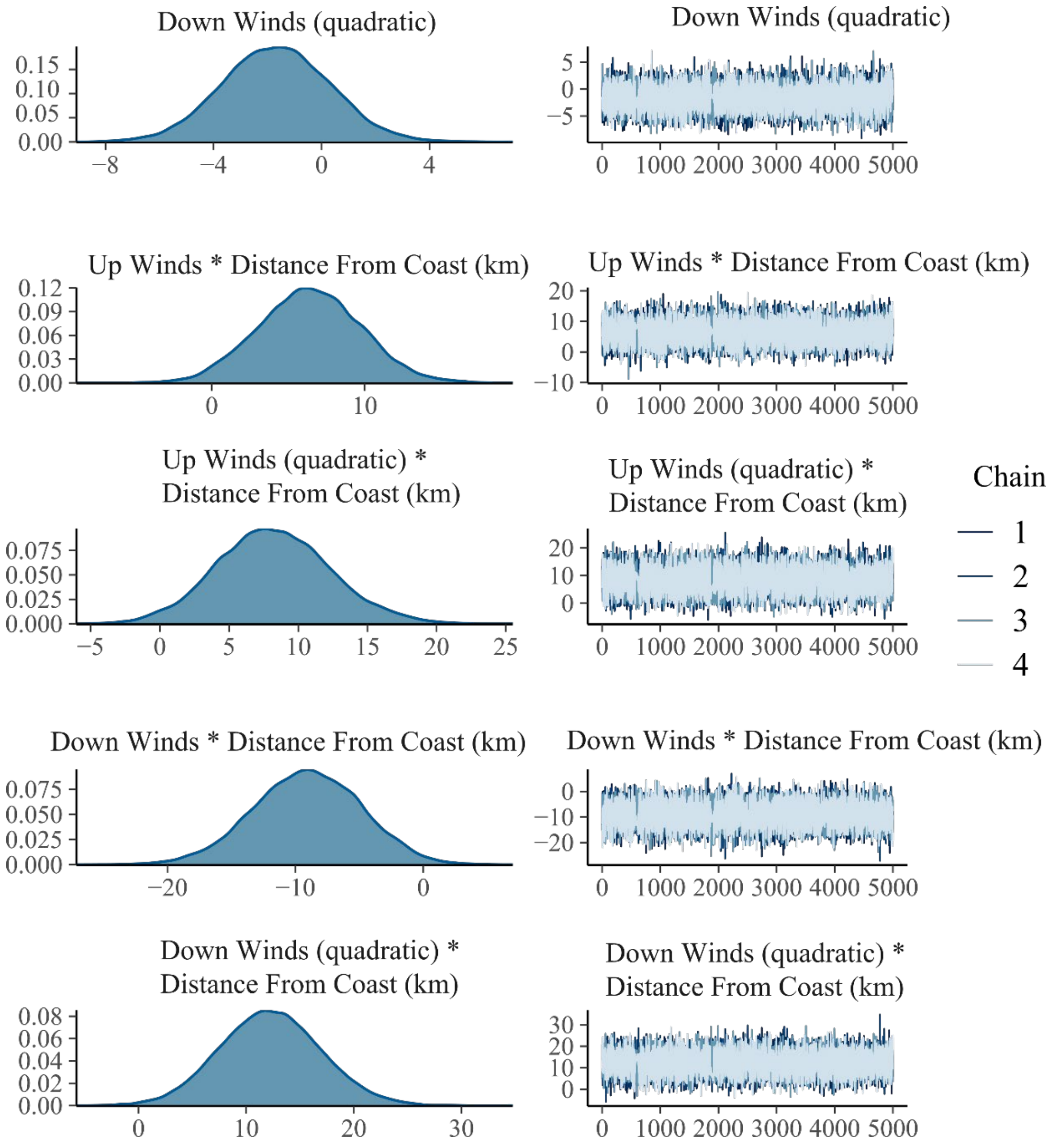


**Figure S7.2** Diagnostic plots of the parameters in the 14 day winds coastal larval fish Bayesian mixed model. Left hand panels show the distribution of the posterior estimates from 10,000 iterations. Right hand panels show trace plots for the 4 chains post warm-up. Continued over the page and on previous page.

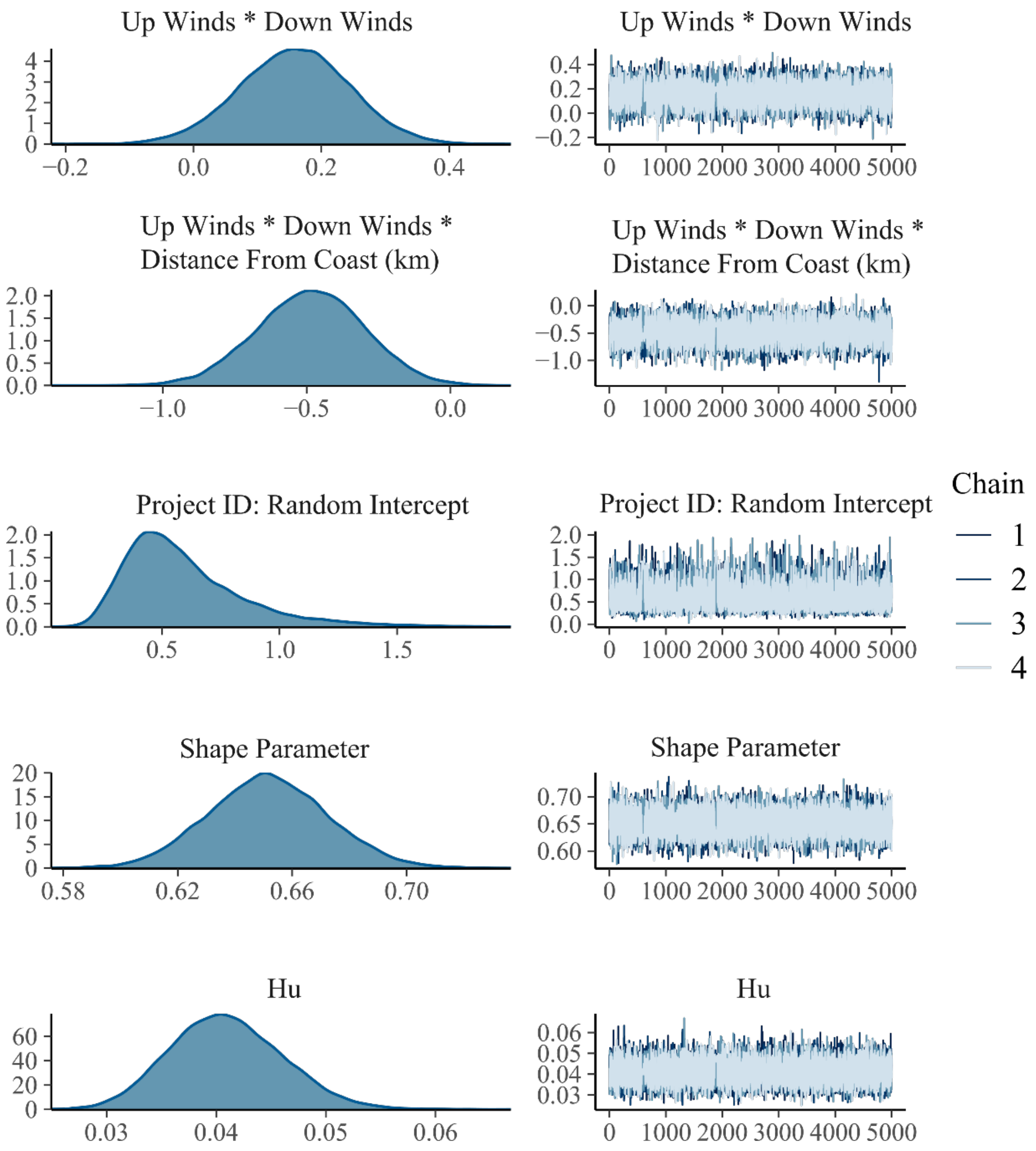


**Figure S7.3** Diagnostic plots of the parameters in the coastal larval fish Bayesian linear mixed model. Left hand panels show the distribution of the posterior estimates from 10,000 iterations. Right hand panels show the trace plots for the 4 chains post warm-up. Continued from previous page.

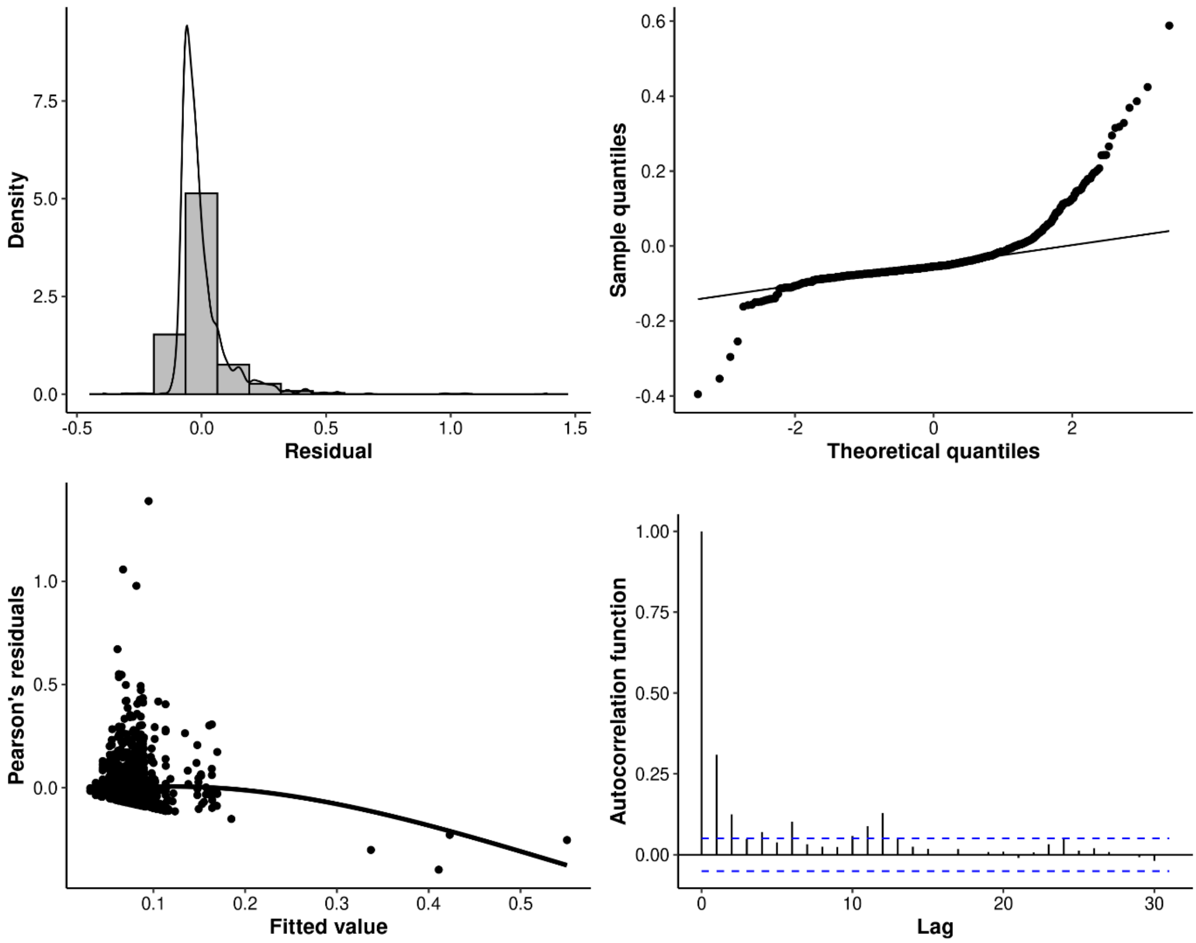


**Figure S8** Diagnostic plots for the coastal larval fish Bayesian linear mixed model. Plots are a density histogram of residuals (top-left), normal quantile-quantile plot (top-right), Pearson's residuals against fitted values (bottom-left) and the autocorrelation function (bottom-right).

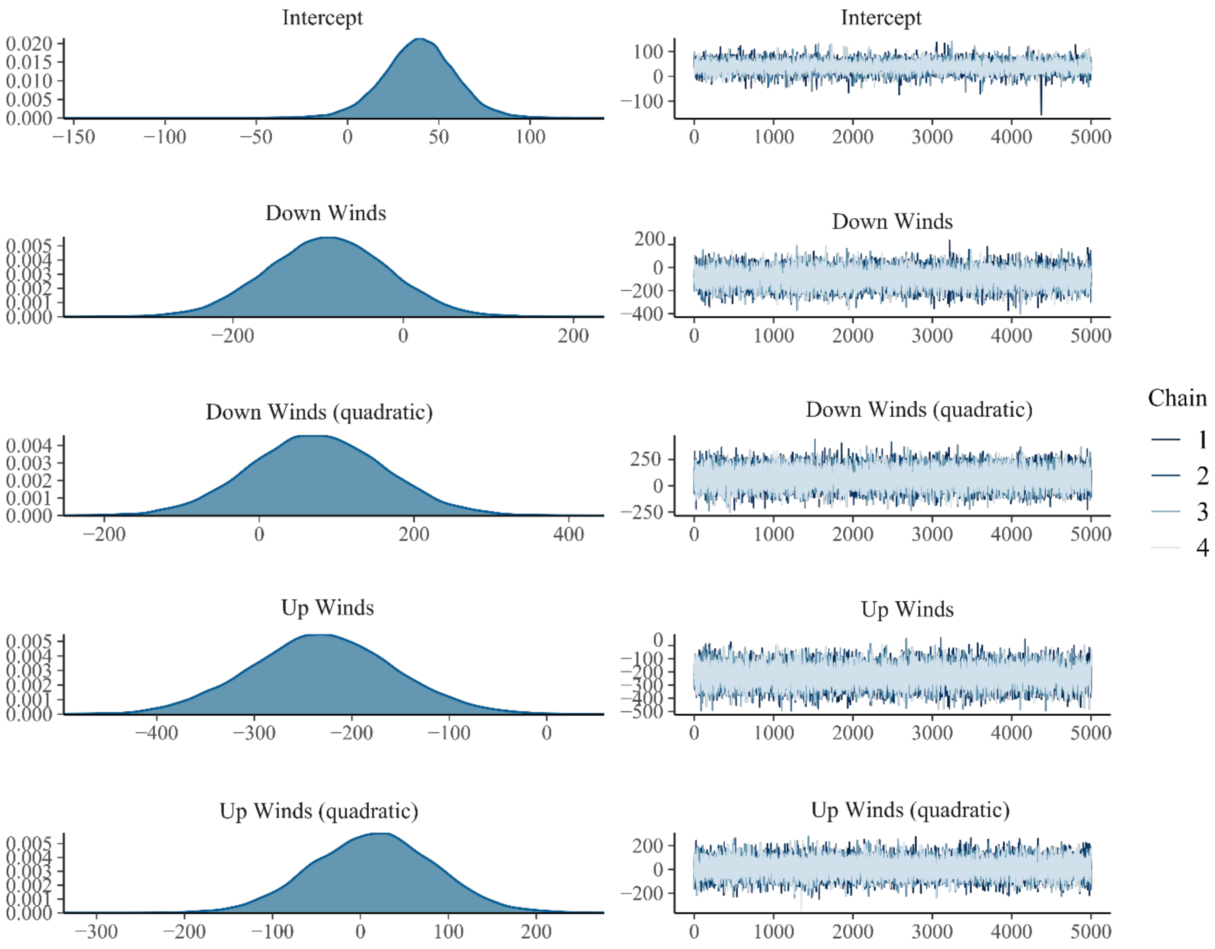


**Figure S9.1** Diagnostic plots of the parameters in the Catch-Per-Unit-Effort (CPUE) Bayesian linear mixed model. Left hand panels show the distribution of the posterior estimates from 20,000 iterations. Right hand panels show trace plots for the 4 chains post warm-up. Continued over the page.

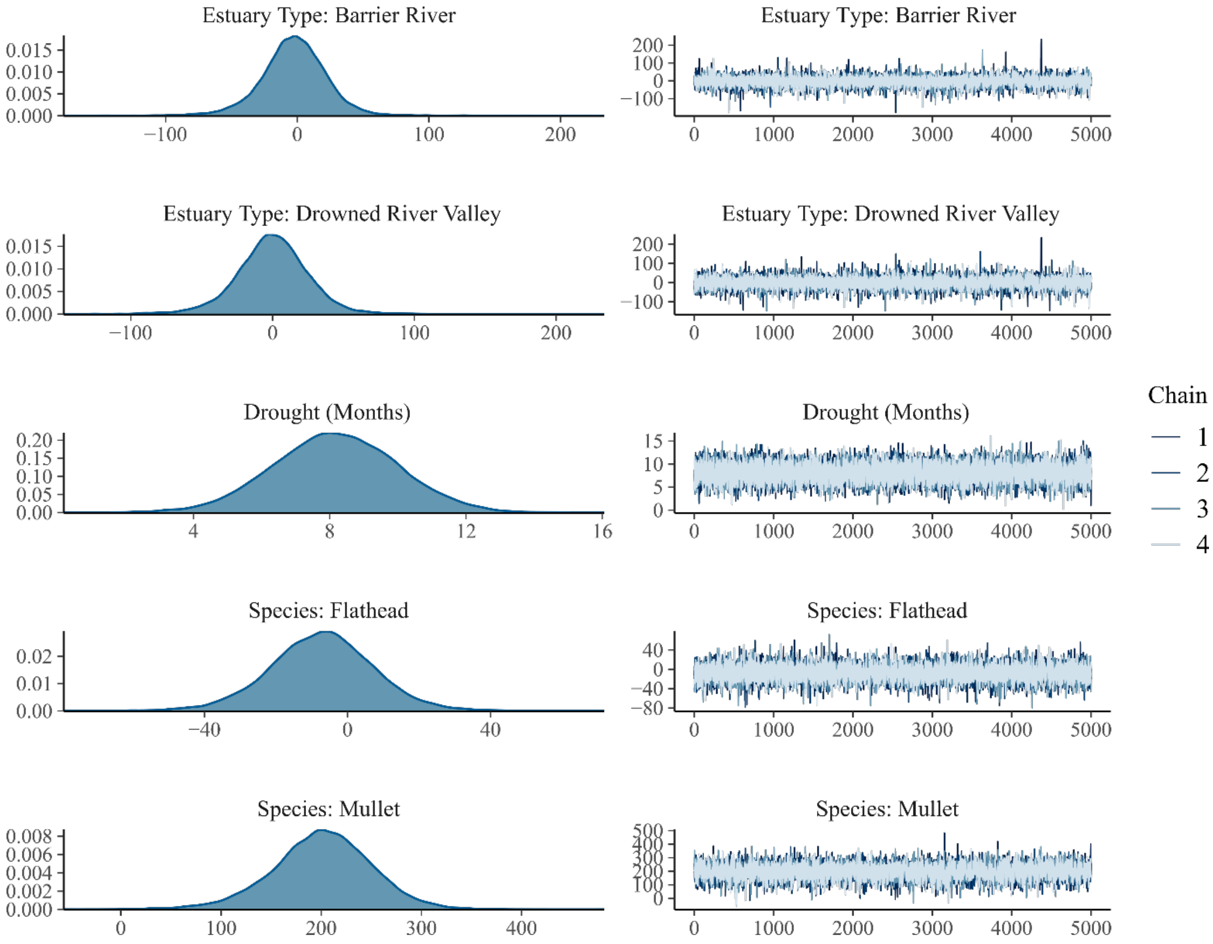


**Figure S9.2** Diagnostic plots of the parameters in the Catch-Per-Unit-Effort (CPUE) Bayesian linear mixed model. Left hand panels show the distribution of the posterior estimates from 20,000 iterations. Right hand panels show trace plots for the 4 chains post warm-up. Continued from previous page and continues over the page.

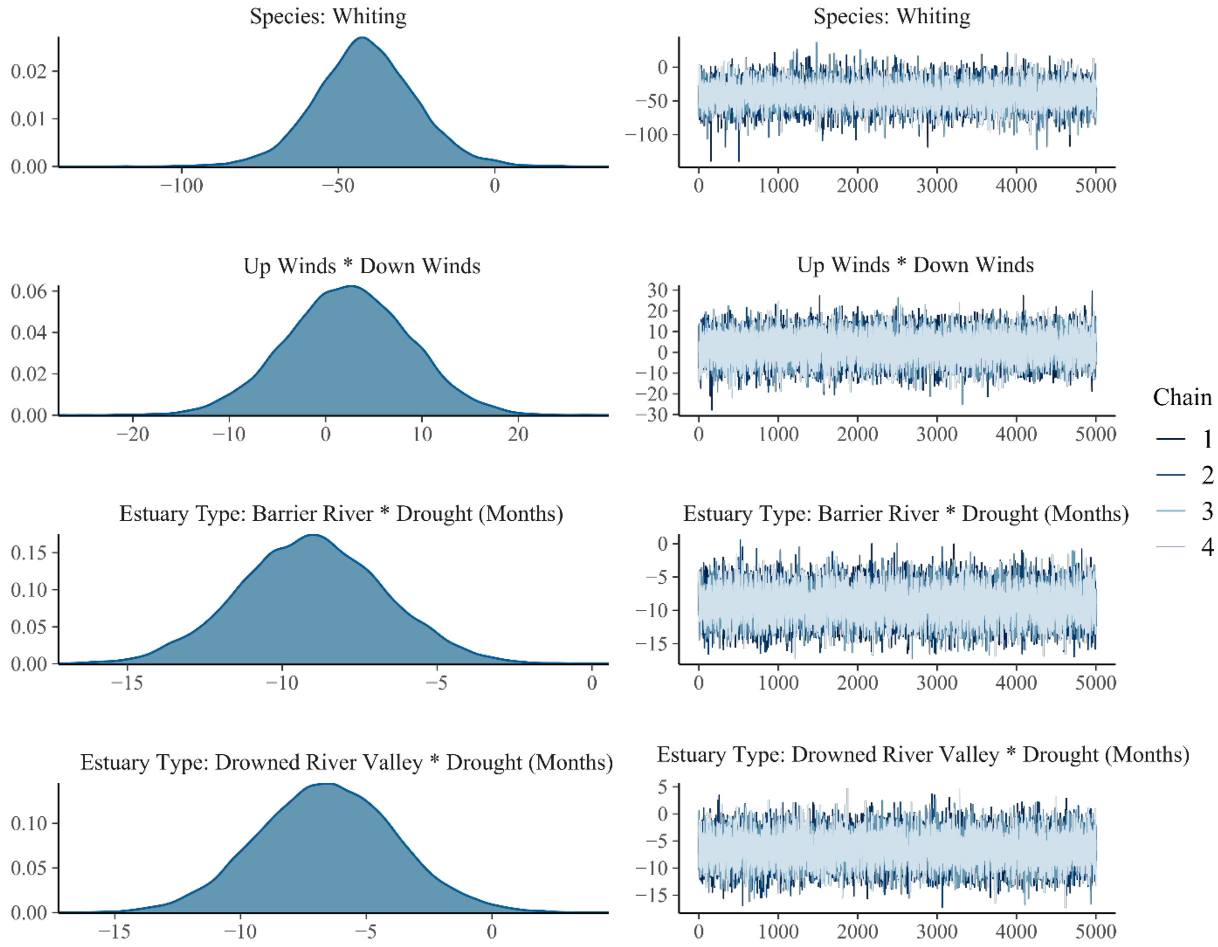


**Figure S9.3** Diagnostic plots of the parameters in the Catch-Per-Unit-Effort (CPUE) Bayesian linear mixed model. Left hand panels show the distribution of the posterior estimates from 20,000 iterations. Right hand panels show trace plots for the 4 chains post warm-up. Continued from previous page.

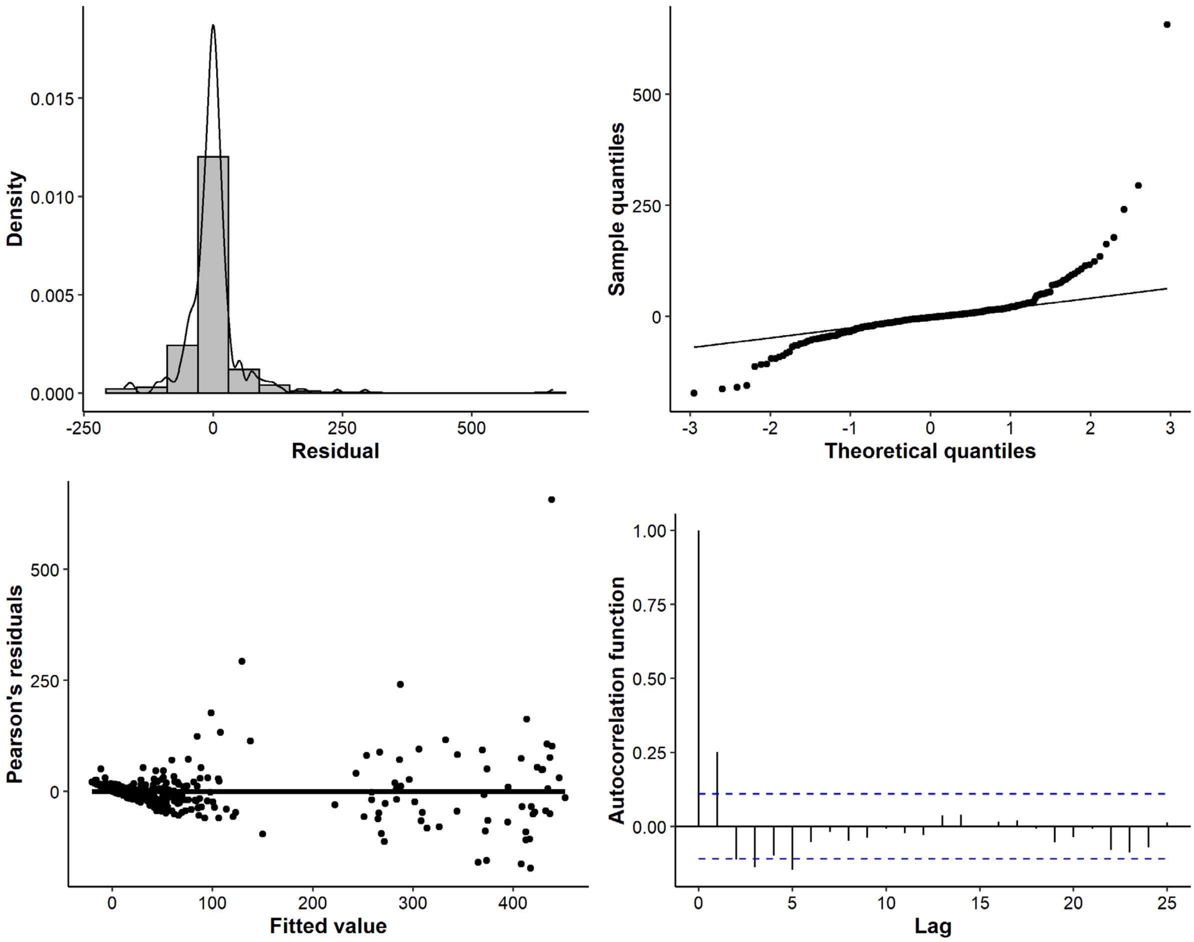


**Figure S10** Diagnostic plots for the Catch-Per-Unit-Effort Bayesian linear mixed model. Plots are a density histogram of residuals (top-left), normal quantile-quantile plot (top-right), Pearson's residuals against fitted values (bottom-left) and the autocorrelation function (bottom-right).

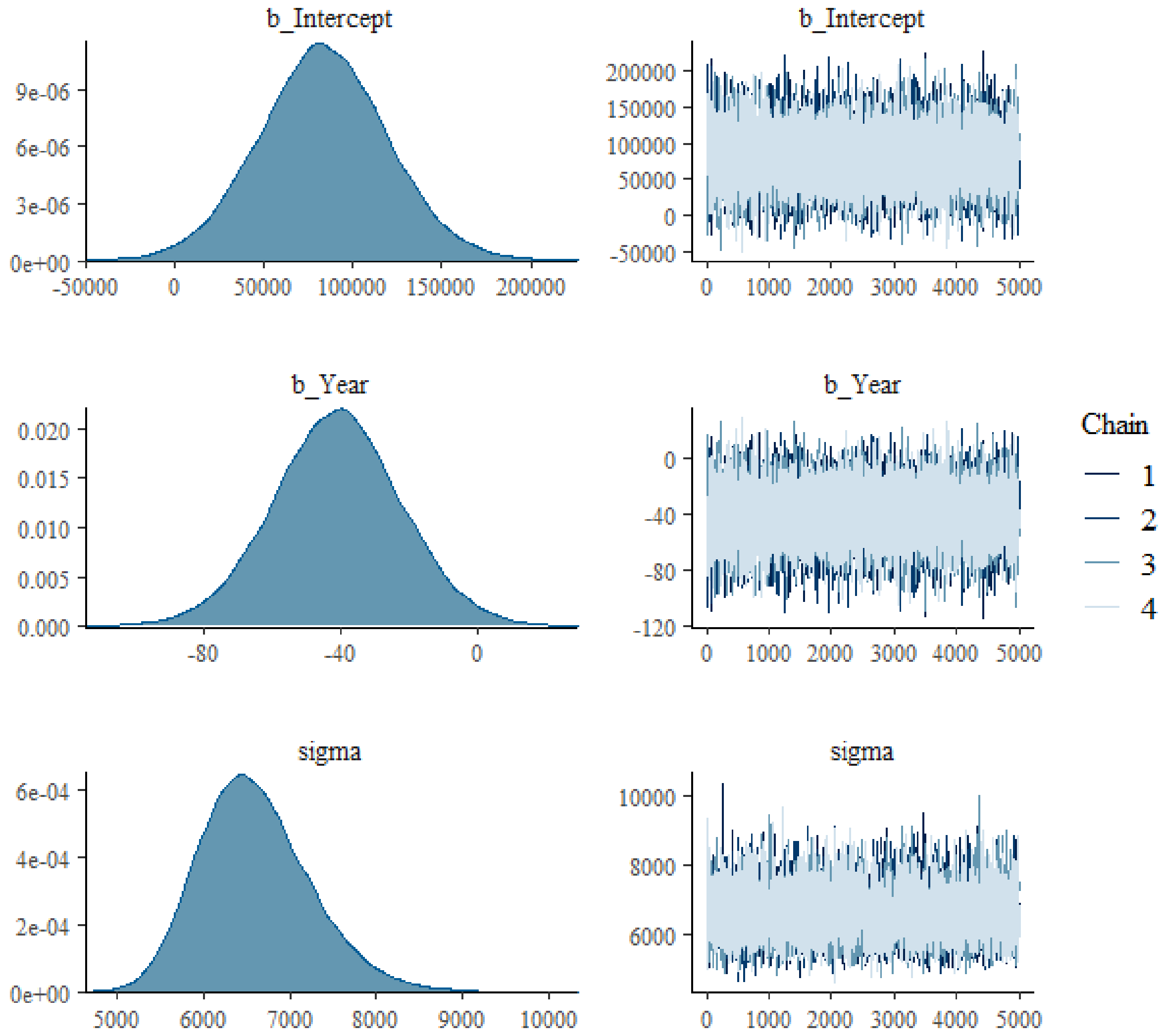


**Figure S11** Diagnostic plots of the parameters in the downwelling favourable (south-easterly) historical wind model. Left hand panels show the posterior distribution of the estimates from 20,000 iterations. Right hand panels show trace plots for the 4 chains post warm-up.

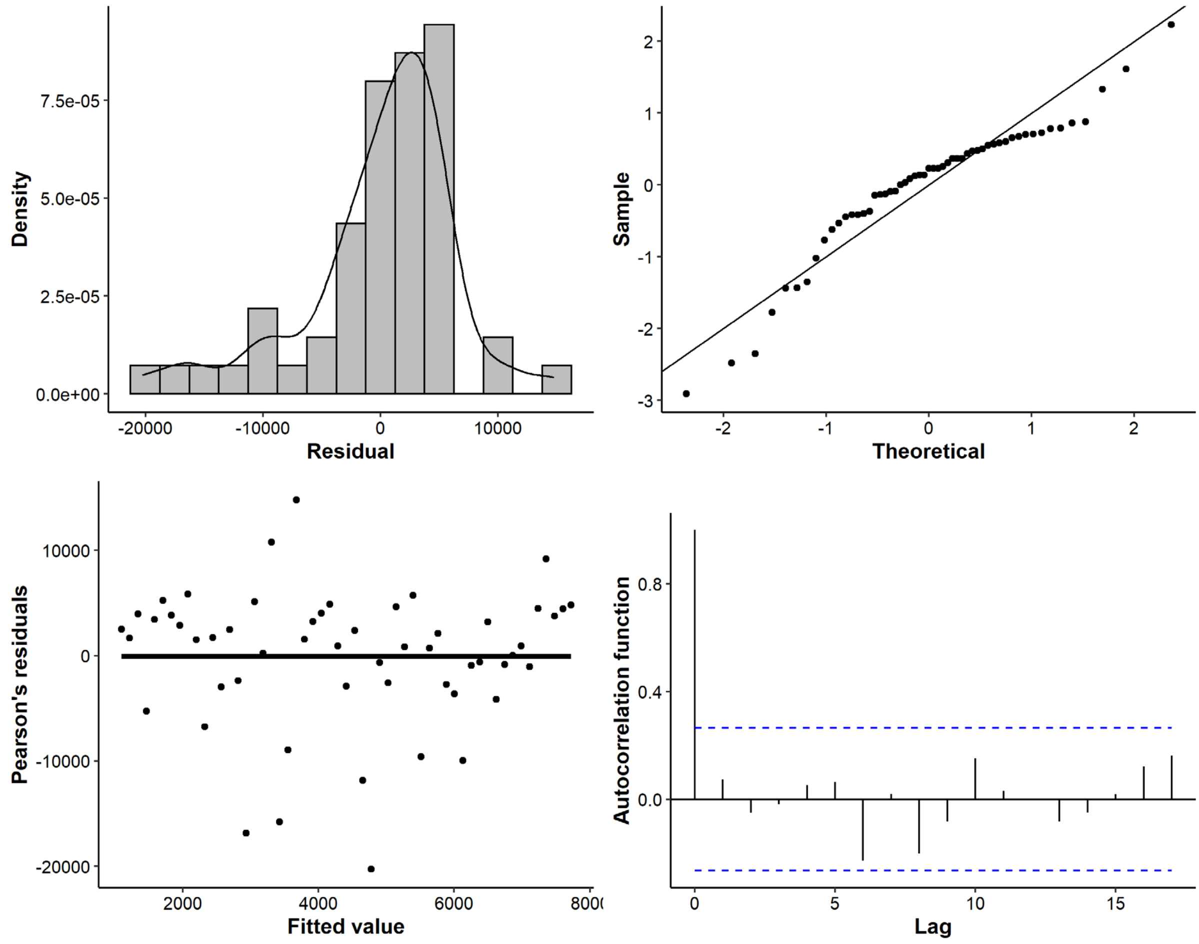


**Figure S12** Diagnostic plots for the downwelling favourable (south-easterly) historical wind model. Plots are a density histogram of residuals (top-left), normal quantile-quantile plot (top-right), Pearson's residuals against fitted values (bottom-left) and the autocorrelation function (bottom-right).

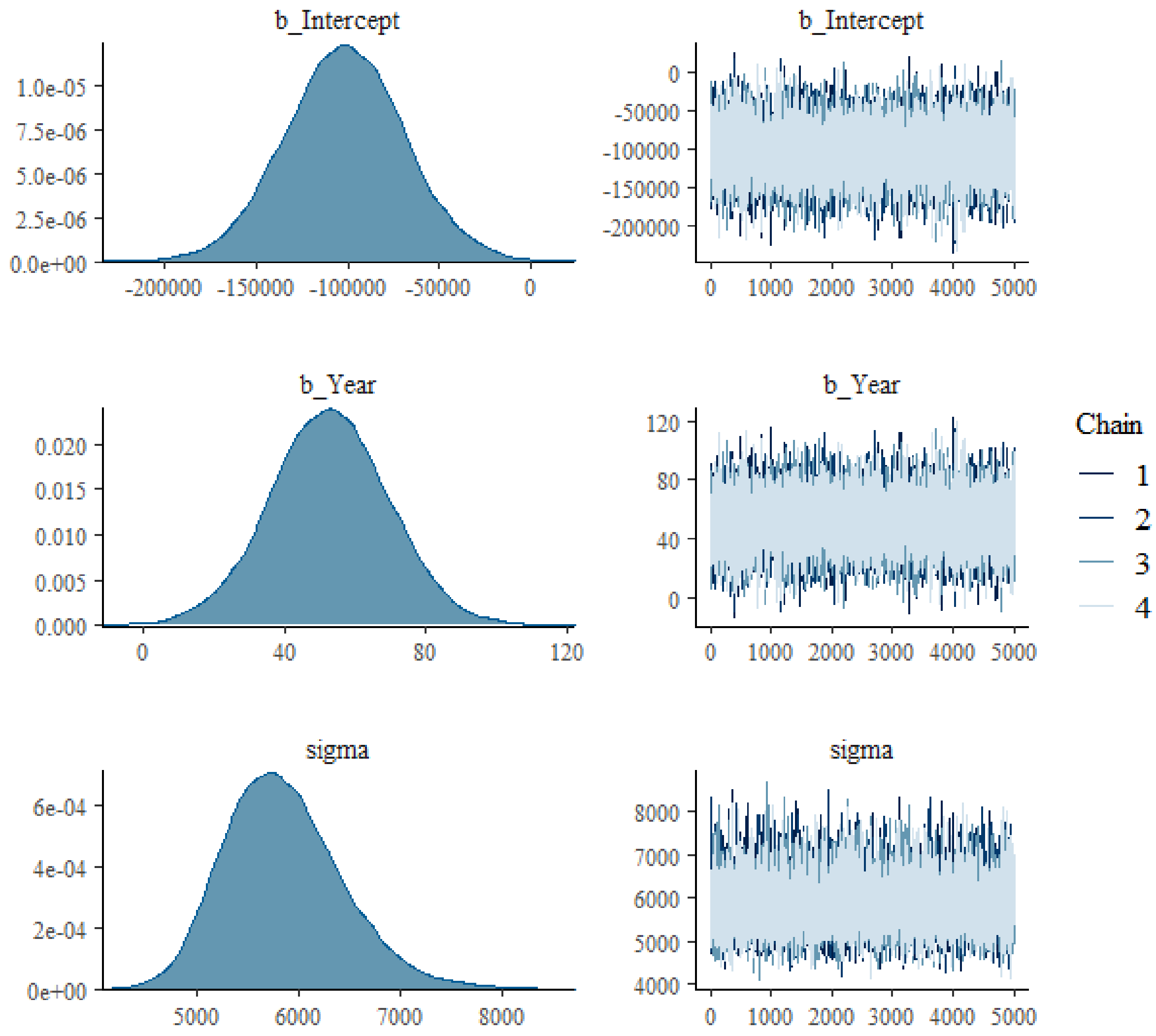


**Figure S13** Diagnostic plots of the parameters in the upwelling favourable (north-easterly) historical wind model. Left hand panels show the distribution of the posterior estimates from 20,000 iterations. Right hand panels show trace plots for the 4 chains post warm-up.

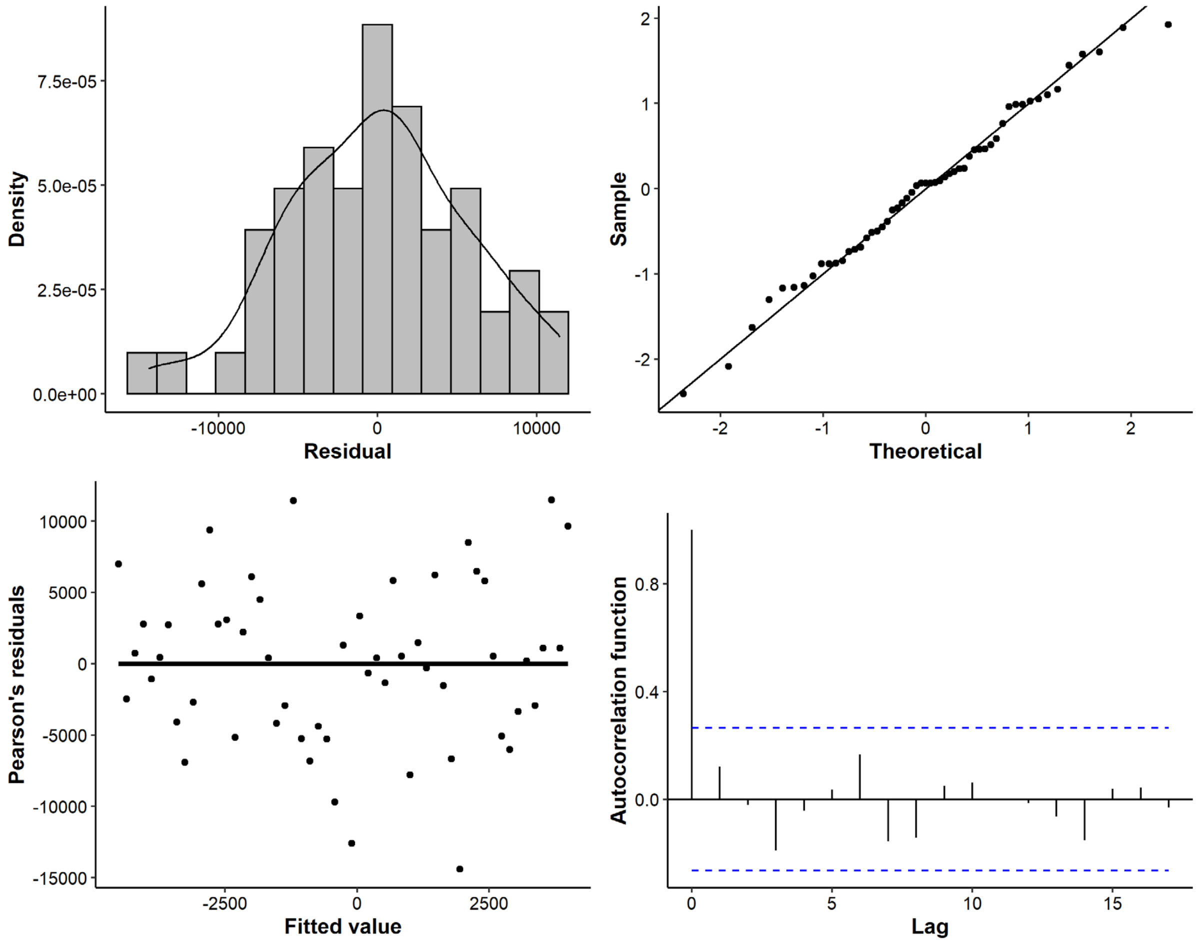


**Figure S14** Diagnostic plots for the upwelling favourable (north-easterly) historical wind model. Plots are a density histogram of residuals (top-left), normal quantile-quantile plot (top-right), Pearson's residuals against fitted values (bottom-left) and the autocorrelation function (bottom-right).

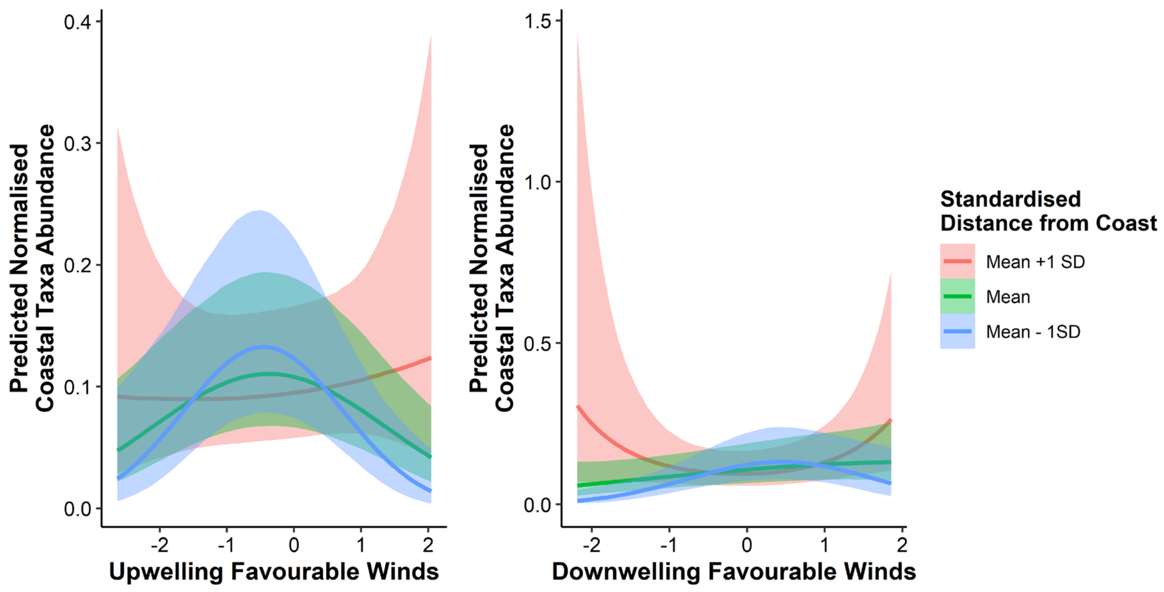


**Figure S15** Visualisation of how the relationships between predicted normalised coastal taxa abundance and upwelling/downwelling favourable winds varies at different distances from the coast.

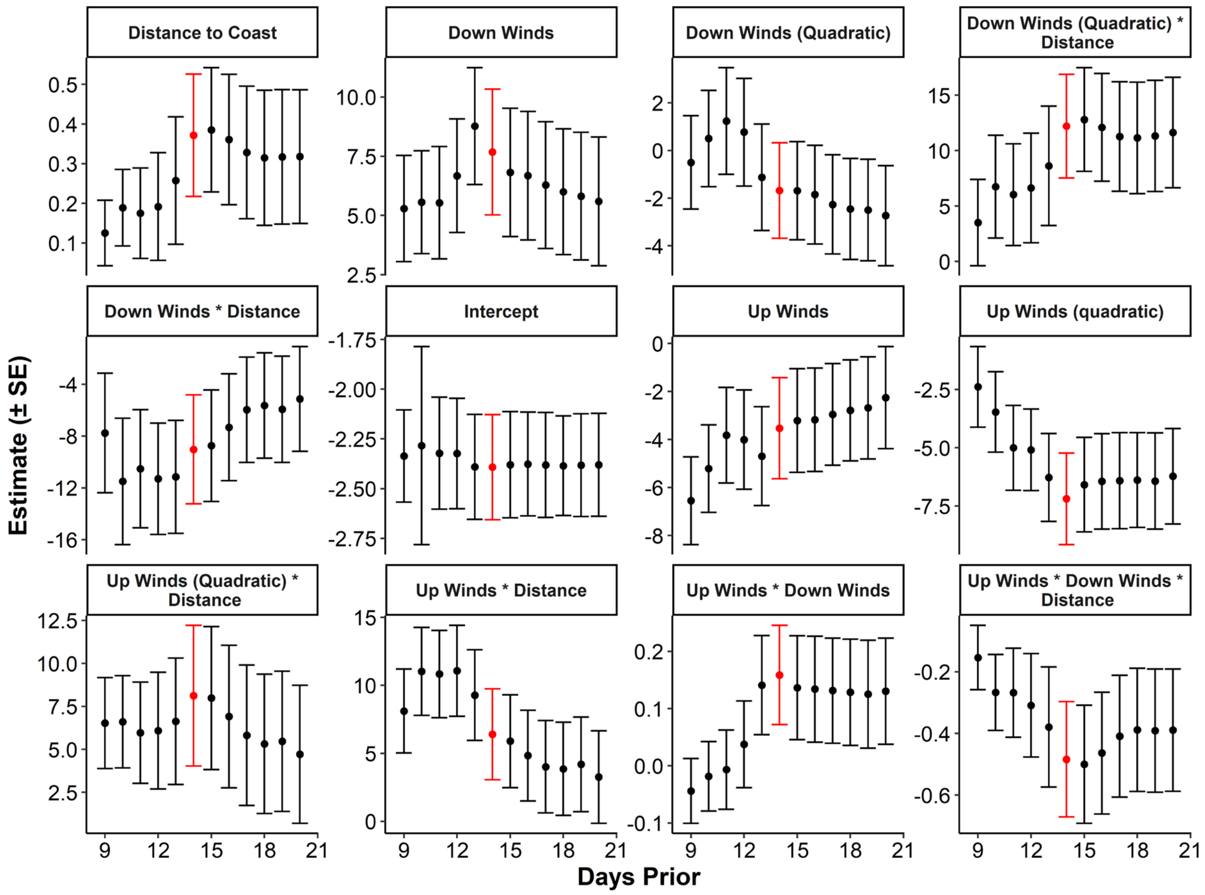


**Figure S16** Results of the sensitivity analysis on the larval fish Bayesian generalised linear mixed model, where the lead up time of the winds was varied (9 – 20 days). Each subplot shows the estimate ± one SE of a model parameter for the various lags. Our main model used a 14 day wind timeframe (here shown in red). We can see the results are generally robust with no abrupt changes in the direction or magnitude of the parameter estimates.

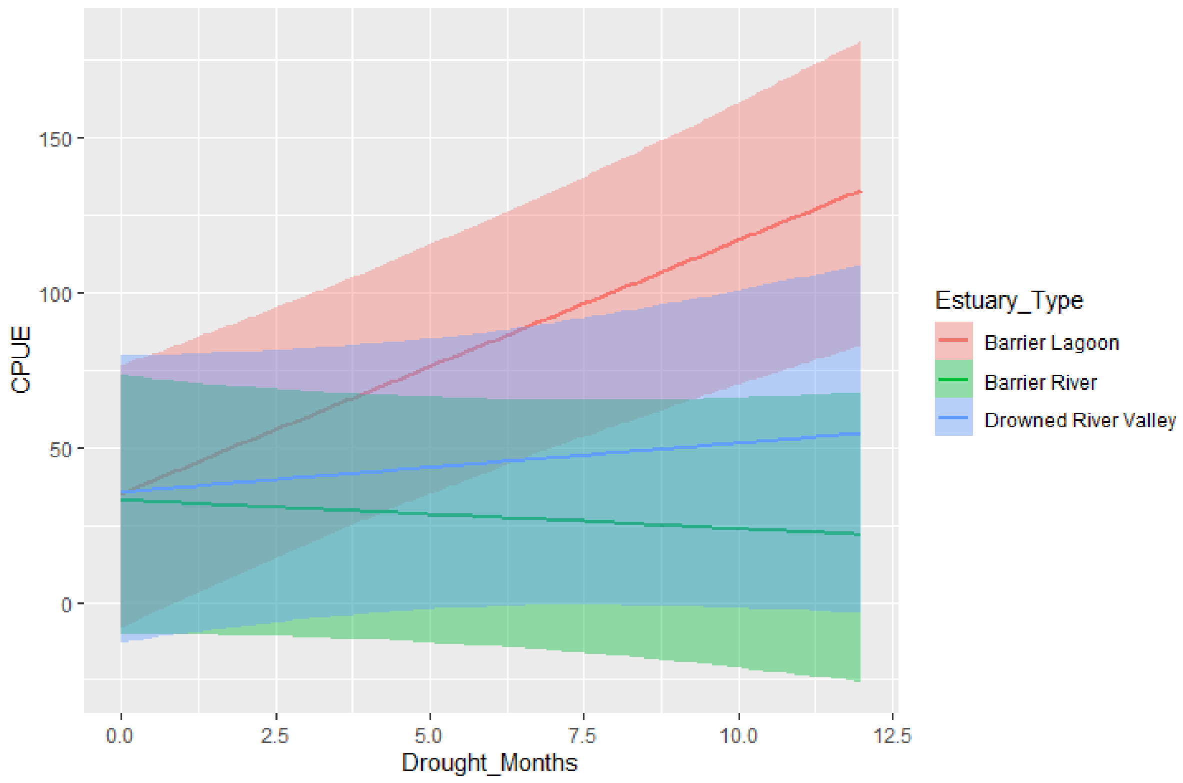


**Figure S17** Predicted effects of increasing drought on annual catch-per-unit-effort (CPUE) based upon the interaction between drought months and estuary type in the multispecies Bayesian linear mixed model.

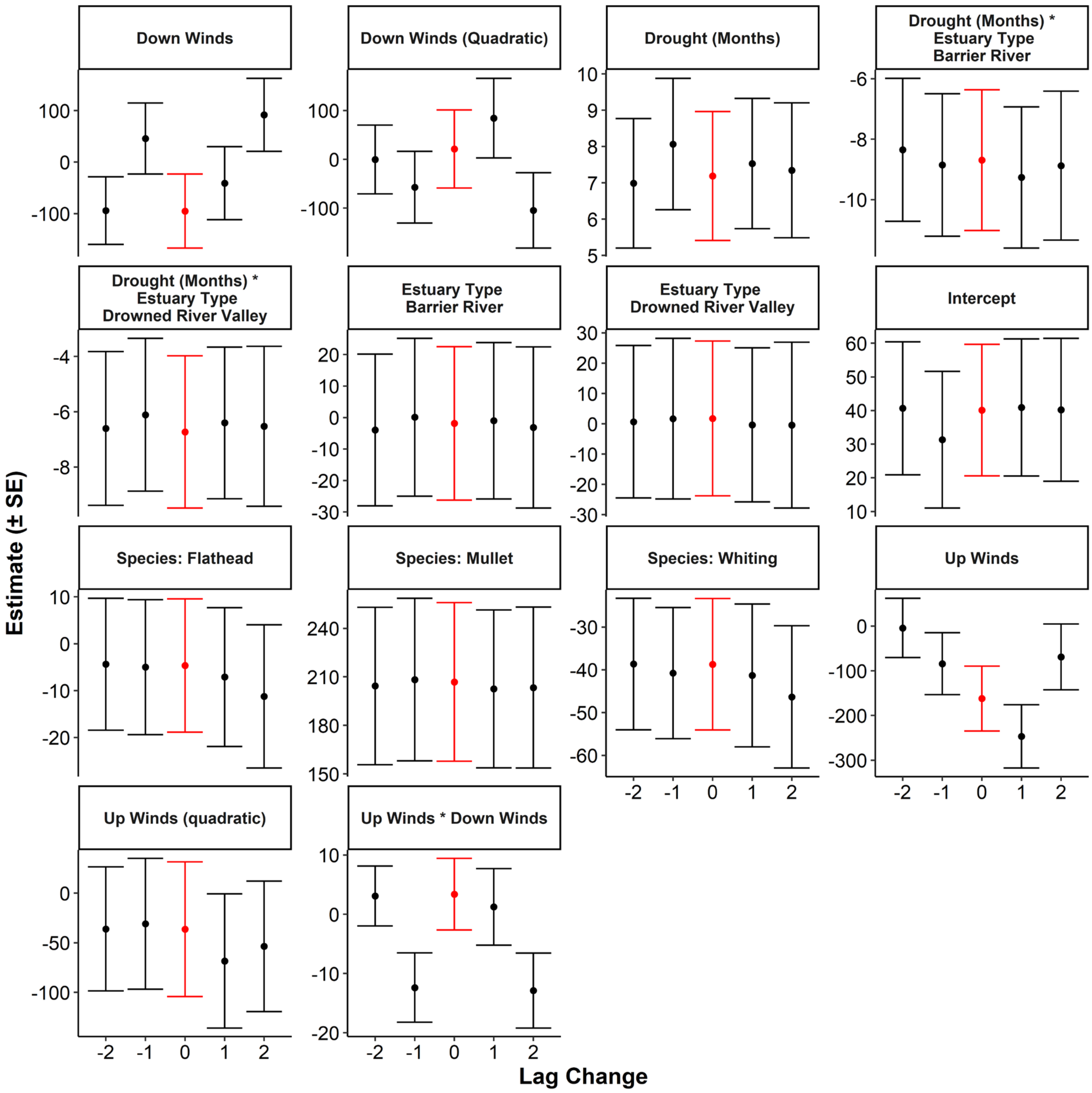


**Figure S18** Results of the sensitivity analysis on the Bayesian linear mixed model where the lag times of the spawning period winds were varied by ±2 years from the chosen lags. Each subplot shows the estimate ± one SE of a model parameter for the various lags. Our main model used lags detailed in Table 1 and this plot shows the effects of changing all of these lags ±2 years from our chosen lags (here shown in red).

Table S1 Sampling Details for the three aims. Note full details of the larval fish data can be found in Smith *et al.* (2018) and further details of the estuarine fisheries dataset can be found in Gillson *et al.* (2009).

| Aim | Grouping Effect (Random Intercept in Model) | Temporal Range (July – June) | Spatial Range (°S) | Number of samples | Other Factor | Note |
|---|---|---|---|---|---|---|
| **1. Larval Abundance** | Project: Franklin_1994 | 1994 | 33.9 - 34 | 223 | Distance from Land (km): 2.53 - 37.9 | |
| | Project: Franklin_1998 | 1998-1999 | 30.6 - 34.1 | 132 | Distance from Land (km): 2.2 - 21.8 | |
| | Project: Investigator_2015 | 2015 | 31.9 - 32.5 | 22 | Distance from Land (km): 16.9 - 28.1 | |
| | Project: Kamala_1989 | 1989-1993 | 33.7 - 34.1 | 1029 | Distance from Land (km): 1.6 - 17.0 | |
| | Project: NIMO_NRS | 2014-2016 | 34.1 | 44 | Distance from Land (km): 2.25 - 5.74 | |
| | Project: Southern_Surveyor_2004 | 2004 | 30.0 - 33.9 | 24 | Distance from Land (km): 6.84 - 53.3 | |
| | Project: Southern_Surveyor_2010 | 2010 | 31.9 - 35.4 | 15 | Distance from Land (km): 11.3 -49.4 | |
| **2. Estuarine Fisheries** | Estuary: Camden River | 1997 - 2007 | 31.6 | 39 | Estuary Type: Barrier River | 4 species |
| | Estuary: Clarence River | 1997 - 2007 | 29.4 | 39 | Estuary Type: Barrier River | 4 species |
| | Estuary: Hawkesbury River | 1997 - 2007 | 33.5 | 39 | Estuary Type: Drowned River Valley | 4 species |
| | Estuary: Hunter River | 1997 - 2007 | 32.9 | 39 | Estuary Type: Barrier River | 4 species |
| | Estuary: Lake Illawarra | 1997 - 2007 | 34.5 | 39 | Estuary Type: Barrier Lagoon | 4 species |

| | | | | | | |
|---|---|---|---|---|---|---|
| | Estuary: Port Stephens | 1997 - 2007 | 32.7 | 39 | Estuary Type: Drowned River Valley | 4 species |
| | Estuary: Tuggerah Lakes | 1997 - 2007 | 33.3 | 39 | Estuary Type: Barrier Lagoon | 4 species |
| | Estuary: Wallis Lake | 1997 - 2007 | 32.2 | 39 | Estuary Type: Barrier Lagoon | 4 species |
| **3a. Historical Winds: Upwelling Favourable** | NA | 1851 – 2014 | 33.7 | 55 | NA | Only used every 3rd year to control autocorrelation |
| **3b. Historical Winds: Downwelling Favourable** | NA | 1851 - 2014 | 33.7 | 55 | NA | Only used every 3rd year to control autocorrelation |

**Table S2** Habitat Classification of the larval fish taxa in the Smith *et al.* (2018) database (NIMO). Larval habitat classification based upon Neira *et al.* (1998) and Miskiewicz (Unpublished Data). The categories were defined as follows: Estuarine - usually spawn in estuaries and juveniles and adults mainly occur in estuaries but can occasionally occur in coastal waters as larvae or adults; Estuarine/coast - taxa in this category spawn in coastal waters and usually recruit to estuaries as larvae/juveniles and the juveniles and adults mainly occur in estuaries but adults can also occur in coastal waters; Coast - taxa in this category spawn in coastal waters and the juveniles and adults also reside and coastal waters on the shelf and are associated with demersal habitats such rocky reefs and soft substrates but can move up in the water column to feed; Coast/pelagic - similar to the coast category but they are usually mobile schooling species but spend most of their time up in the water column rather than associating with the seafloor; Coast/slope - similar to the coast category but the distribution of adults can extend further offshore over the shelf and onto the slope; Tropical – Species generally spawned in tropical waters; Ocean – usually found in deep water ocean as both larvae and adults; Slope: adults and larvae usually found on the slope of the continental shelf. The larval fish analysis in the current paper combined the Coast, Coast/pelagic, Coast/slope and Estuarine/coast categories as "coastal".

| Category | Family-NIMO | Common-NIMO | Species-NIMO | NIMO_NAME |
|---|---|---|---|---|
| Tropical | Acanthuridae | Surgeonfish | | Acanthuridae_37437000 |
| Coast/slope | Acropomatidae | Threespine cardinalfish | *Apogonops anomalus* | Acropomatidae_Apogonops.anomalus_37311053 |
| Coast/slope | Acropomatidae | Lanternbelly | *Synagrops* spp | Acropomatidae_Synagrops.spp_37311949 |
| Tropical | Ammodytidae | Sand lance | | Acropomatidae_other_37311956 |
| Tropical | Anguilliformes | Order of eels | | Ammodytidae_37425000 |
| Coast/slope | Antennariidae | Anglerfish | | Anguilliformes_37990019 |
| Coast | Anthiinae | Sea perch | | Anthiinae_37311907 |
| Tropical | Aploactinidae | Velvetfish | *Matsubarichthys inusitatus* | Aploactinidae_Matsubarichthys.inusitatus_37290013 |
| Coast | Aplodactylidae | Sea carp | *Aplodactylus* spp | Aplodactylidae_Aplodactylus.spp_37376901 |
| Coast | Apogonidae | Cardinal fish | | Apogonidae_37327926 |
| Ocean | Argentinidae | Herring smelt | | Argentinidae_37097000 |
| Coast/pelagic | Arripidae | Australian salmon | *Arripis trutta* | Arripidae_Arripis.trutta_37344002 |
| Ocean | Astronesthidae | Dragonfish | | Astronesthidae_37108000 |

| Category | Family-NIMO | Common-NIMO | Species-NIMO | NIMO_NAME |
|---|---|---|---|---|
| Estuary | Atherinidae | Hardyhead | | Atherinidae_37246000 |
| Coast | Aulopidae | Sergeant Baker | *Hime* spp | Aulopidae_Hime.spp_37117902 |
| Tropical | Balistidae | Triggerfish | | Balistidae_37465900 |
| Ocean | Bathylagidae | Deepsea smelt | | Bathylagidae_37098000 |
| Coast | Berycidae | Redfish | *Centroberyx affinis* | Berycidae_Centroberyx.affinis_37258003 |
| Coast | Berycidae | Redfish | *Beryx* spp | Berycidae_Beryx.spp_37258901 |
| Estuary/coast | Blenniidae | Blenny | *Petroscirtes lupus* | Blenniidae_Petroscirtes.lupus_37408073 |
| Estuary/coast | Blenniidae | Blenny | *Omobranchus anolius* | Blenniidae_Omobranchus.anolius_37408058 |
| Estuary/coast | Blenniidae | Blenny | *Parablennius* spp | Blenniidae_Parablennius.spp_37408918 |
| Tropical | Blenniidae | Blenny | *Plagiotremus* spp | Blenniidae_Plagiotremus.spp_37408921 |
| Coast | Bothidae | Flatfish | *Lophonectes gallus* | Bothidae_Lophonectes.gallus_37460001 |
| Tropical | Bothidae | Flatfish | *Crossorhombus* spp | Bothidae_Crossorhombus.spp_37460907 |
| Tropical | Bothidae | Flatfish | *Engyprosopon* spp | Bothidae_Engyprosopon.spp_37460908 |
| Tropical | Bothidae | Flatfish | *Grammatobothus* spp | Bothidae_Grammatobothus.spp_37460909 |
| Coast | Bothidae | Flatfish | *Arnoglossus* spp | Bothidae_Arnoglossus.spp_37460902 |
| Tropical | Bothidae | Flatfish | *Asterorhombus* spp | Bothidae_Asterorhombus.spp_37460903 |
| Coast | Bovichtidae | Thornfishes | *Bovichtus augustifrons* | Bovichtidae_Bovichtus.augustifrons_37403001 |
| Coast | Bovichtidae | Thornfishes | *Pseudaphritis urvilli* | Bovichtidae_Pseudaphritis.urvilli_37403003 |
| Coast/pelagic | Bramidae | Pomfret | *Brama* spp | Bramidae_Brama.spp_37342900 |

| Category | Family-NIMO | Common-NIMO | Species-NIMO | NIMO_NAME |
|---|---|---|---|---|
| Coast/slope | Bregmacerotidae | Codlets | *Bregmaceros* spp | Bregmacerotidae_Bregmaceros.spp_37225901 |
| Coast | Callanthiidae | Splendid perch | *Callanthias australis* | Callanthiidae_Callanthias.australis_37311055 |
| Coast | Callionymidae | Dragonets | | Callionymidae_37427917 |
| Tropical | Caproidae | Boarfishes | *Antigonia* spp | Caproidae_Antigonia.spp_37267901 |
| Coast/pelagic | Carangidae | Scad | *Pseudocaranx georgianus* | Carangidae_Pseudocaranx.georgianus_37337062 |
| Coast/pelagic | Carangidae | Jack mackerel | *Trachurus declivus* | Carangidae_Trachurus.declivus_37337002 |
| Coast/pelagic | Carangidae | Scad | *Decapterus* spp | Carangidae_Decapterus.spp_37337901 |
| Coast/pelagic | Carangidae | Yellowtail scad | *Trachurus novaezelandiae* | Carangidae_Trachurus.novaezelandiae_37337003 |
| Coast/pelagic | Carangidae | Kingfish | *Seriola* spp | Carangidae_Seriola.spp_37337927 |
| Tropical | Carapidae | Pearlfish | | Carapidae_37229000 |
| Coast/slope | Centrolophidae | Trevallas | *Seriolella* spp | Centrolophidae_Seriolella.spp_37445909 |
| Coast | Cepolidae | Bandfish | *Acanthocepola* spp | Cepolidae_Acanthocepola.spp_37380901 |
| Coast | Cepolidae | Bandfish | *Owstonia* spp | Cepolidae_Owstonia.spp_37380903 |
| Coast | Cepolidae | Bandfish | *Cepola australis* | Cepolidae_Cepola.australis_37380001 |
| Ocean | Cetomimidae | Whalefish | | Cetomimidae_37132000 |
| Tropical | Chaetodontidae | Butterfly fish | | Chaetodontidae_37365900 |
| Tropical | Champsodontidae | Gapers | | Champsodontidae_37401000 |
| Estuary/coast | Chandidae | Port Jackson perchlet | *Ambassis jacksoniensis* | Chandidae_Ambassis.jacksoniensis_37310012 |
| Estuary | Chandidae | Estuary perchlet | *Ambassis marianus* | Chandidae_Ambassis.marianus_37310018 |
| Ocean | Chauliodontidae | Viperfish | | Chauliodontidae_37111000 |

| Category | Family-NIMO | Common-NIMO | Species-NIMO | NIMO_NAME |
|---|---|---|---|---|
| Coast | Cheilodactylidae | Jackass morwong | *Nemadactylus macropterus* | Cheilodactylidae_Nemadactylus.macropterus_3737700[illegible] |
| Coast | Cheilodactylidae | Morwong | *Cheilodactylus* spp | Cheilodactylidae_Cheilodactylus.spp_37377902 |
| Ocean | Chiasmodontidae | Swallowers | | Chiasmodontidae_37402000 |
| Coast | Chironemidae | Marblefish | *Chironemus* spp | Chironemidae_Chironemus.spp_37375901 |
| Coast/slope | Chlorophthalmidae | Greeneyes | *Chlorophthalmus* spp | Chlorophthalmidae_Chlorophthalmus.spp_37120903 |
| Tropical | Cirrhitidae | Hawkfish | | Cirrhitidae_37374000 |
| Estuary/coast | Clinidae | Weedfish | | Clinidae_37416000 |
| Coast/pelagic | Clupeidae | Pilchard | *Sardinops sagax* | Clupeidae_Sardinops.sagax_37085002 |
| Coast/pelagic | Clupeidae | Maray | *Etrumeus teres* | Clupeidae_Etrumeus.teres_37085001 |
| Coast/pelagic | Clupeidae | Blue sprat | *Spartelloides robustus* | Clupeidae_Spartelloides.robustus_37085003 |
| Coast/pelagic | Clupeidae | Sandy sprat | *Hyperlophus vittatus* | Clupeidae_Hyperlophus.vittatus_37085005 |
| Tropical | Coryphaenidae | Dolphin fish | *Coryphena* spp | Coryphaenidae_Coryphena.spp_37338901 |
| Coast | Creediidae | Sand burrower | *Creedia* spp | Creediidae_Creedia.spp_37395901 |
| Coast | Creediidae | Sand burrower | *Limnichthys* spp | Creediidae_Limnichthys.spp_37395902 |
| Coast | Cynoglossidae | Tongue sole | | Cynoglossidae_37463000 |
| Tropical | Dactylopteridae | Flying gurnard | *Dactyloptena* spp | Dactylopteridae_Dactyloptena.spp_37308902 |
| Coast | Dinolestidae | Longfin pike | *Dinolestis lewini* | Dinolestidae_Dinolestis.lewini_37327002 |
| Ocean | Diretmidae | Discfish | *Diretmus* spp | Diretmidae_Diretmus.spp_37254902 |
| Coast/pelagic | Emmelichthys | Redbait | *Emmelichthys nitidus* | Emmelichthys_Emmelichthys.nitidus_37345001 |

| Category | Family-NIMO | Common-NIMO | Species-NIMO | NIMO_NAME |
|---|---|---|---|---|
| Coast/pelagic | Engraulidae | Anchovy | *Engraulis australis* | Engraulidae_Engraulis.australis_37086001 |
| Coast | Enoplosidae | Old wife | *Enoplosus armatus* | Enoplosidae_Enoplosus.armatus_37366001 |
| Tropical | Epinephelinae | Grouper | | Epinephelinae_37311908 |
| Ocean | Evermannellidae | Sabretooth | | Evermannellidae_37130000 |
| Tropical | Exocoetidae | Flying fish | | Exocoetidae_37233000 |
| Tropical | Fistulariidae | Flutemouth | | Fistulariidae_37278000 |
| Coast | Gempylidae | Barracouda | *Thyrsites atun* | Gempylidae_Thyrsites.atun_37439001 |
| Slope | Gempylidae | Gemfish | *Rexea* solandri | Gempylidae_Rexea.solandri_37439002 |
| Slope | Gempylidae | Snake mackeral | *Gempylus serpens* | Gempylidae_Gempylus.serpens_37439010 |
| Estuary/coast | Gerreidae | Silver belly | *Gerres subfasciatus* | Gerreidae_Gerres.subfasciatus_37349005 |
| Estuary/coast | Gerreidae | Silver belly | *Parequula melbournensis* | Gerreidae_Parequula.melbournensis_37349001 |
| Estuary/coast | Girellidae | Luderick | *Girella tricuspidata* | Girellidae_Girella.tricuspidata_37361007 |
| Estuary/coast | Girellidae | Luderick | *Girella* spp | Girellidae_Girella.spp_37361902 |
| Estuary/coast | Gobiesocidae | Clingfish | *Alabes* spp | Gobiesocidae_Alabes.spp_37206901 |
| Estuary/coast | Gobiesocidae | Clingfish | other | Gobiesocidae_other_37206000 |
| Estuary/coast | Gobiidae | Goby | | Gobiidae_37428000 |
| Coast | Gonorhynchidae | Beak salmon | *Gonorhynchus* greyi | Gonorhynchidae_Gonorhynchus.greyi_37141001 |
| Ocean | Gonostomatidae | Bristlemouth | | Gonostomatidae_37106912 |
| Tropical | Grammistidae | Soapfish | | Grammistidae_37312000 |
| Tropical | Haemulidae | Sweetlips | | Haemulidae_37350000 |

| Category | Family-NIMO | Common-NIMO | Species-NIMO | NIMO_NAME |
|---|---|---|---|---|
| Coast/pelagic | Hemiramphidae | Garfish | *Hemiramphus* spp | Hemiramphidae_Hemiramphus.spp_37234903 |
| Tropical | Holocentridae | Squirelfish | | Holocentridae_37261000 |
| Coast/slope | Hoplichthyidae | Ghost flathead | *Hoplichthys* spp | Hoplichthyidae_Hoplichthys.spp_37297901 |
| Coast/slope | Howellidae | Pelagic bass | *Howella* spp | Howellidae_Howella.spp_37327912 |
| Ocean | Idiacanthidae | Black dragonfish | *Idiacanthus* spp | Idiacanthidae_Idiacanthus.spp_37113901 |
| Ocean | Ipnopidae | Tripodfish | | Ipnopidae_37123000 |
| Tropical | Istiophoridae | Marlin | | Istiophoridae_37444000 |
| Coast | Kyphosidae | Drummer | *Kyphosus* spp | Kyphosidae_Kyphosus.spp_37361903 |
| Estuary/coast | Labridae | Wrasse | | Labridae_37384000 |
| Coast/slope | Lampridiformes | Ribbonfish | | Lampridiformes_37990051 |
| Coast | Latridae | Trumpeter | *Latris lineata* | Latridae_Latris.lineata_37378001 |
| Tropical | Leiognathidae | Ponyfish | | Leiognathidae_37341000 |
| Tropical | Leptobramidae | Beach salmon | | Leptobramidae_37357905 |
| Coast | Leptoscopidae | Sandfish | | Leptoscopidae_37398000 |
| Tropical | Lethrinidae | Emperor | *Lethrinus* spp | Lethrinidae_Lethrinus.spp_37351902 |
| Coast/slope | Lophiiformes | Anglerfish | | Lophiiformes_37990052 |
| Tropical | Lutjanidae | Snapper | | Lutjanidae_37346000 |
| Coast/slope | Macroramphosidae | Bellowsfish | *Macroramphosus* spp | Macroramphosidae_Macroramphosus.spp_37279902 |
| Ocean | Macrouridae | Rattail | | Macrouridae_37232000 |
| Coast | Malacanthidae | Tilefish | *Branchiostegus* spp | Malacanthidae_Branchiostegus.spp_37331901 |
| Coast | Malacanthidae | Blanquillo | *Malacanthus* spp | Malacanthidae_Malacanthus.spp_37331903 |
| Ocean | Melamphaidae | Bigscale | | Melamphaidae_37251000 |

| Category | Family-NIMO | Common-NIMO | Species-NIMO | NIMO_NAME |
|---|---|---|---|---|
| Ocean | Melanostomiidae | Black dragonfish | | Melanostomiidae_37109000 |
| Coast | Microcanthidae | Mado | *Atypichthys strigatus* | Microcanthidae_Atypichthys.strigatus_37361010 |
| Coast | Microcanthidae | Stripey | *Microcanthus strigatus* | Microcanthidae_Microcanthus.strigatus_37361005 |
| Tropical | Microdesmidae | Wormfish | | Microdesmidae_37435000 |
| Ocean | Molidae | Sunfish | | Molidae_37470000 |
| Estuary/coast | Monacanthidae | Leatherjacket | | Monacanthidae_37465903 |
| Coast | Monodactylidae | Pomfred | *Schuetta* spp | Monodactylidae_Schuetta.spp_37356902 |
| Estuary/coast | Monodactylidae | Diamondfish | *Monodactylus argenteus* | Monodactylidae_Monodactylus.argenteus_37356002 |
| Coast | Moridae | Beardies | | Moridae_37224000 |
| Estuary/coast | Mugilidae | Mullet | *Liza argentea* | Mugilidae_Liza.argentea_37381004 |
| Estuary/coast | Mugilidae | Mullet | other | Mugilidae_other_37381000 |
| Tropical | Mullidae | Goatfish | | Mullidae_37355000 |
| Ocean | Myctophidae | Lanternfish | | Myctophidae_37122000 |
| Tropical | Nemipteridae | Threadfin bream | | Nemipteridae_37347000 |
| Coast/slope | Nomeidae | Driftfish | | Nomeidae_37446000 |
| Ocean | Notosudidae | Paperbones | | Notosudidae_37125000 |
| Coast | Odacidae | Rock cale | | Odacidae_37385000 |
| Coast/slope | Ophidiidae | Ling | *Genyptrus* spp | Ophidiidae_Genyptrus.spp_37228901 |
| Tropical | Ophidiidae | Ling | *Brotula* spp | Ophidiidae_Brotula.spp_37228912 |
| Tropical | Ostraciidae | Cowfish | | Ostraciidae_37466000 |
| Ocean | Paralepididae | Barracudinas | | Paralepididae_37126000 |
| Estuary/coast | Paralichthyidae | Large tooth flounder | *Pseudorhombus* spp | Paralichthyidae_Pseudorhombus.spp_37460919 |
| Tropical | Pegasidae | Sea moth | *Pegasus* spp | Pegasidae_Pegasus.spp_37309902 |

| Category | Family-NIMO | Common-NIMO | Species-NIMO | NIMO_NAME |
|---|---|---|---|---|
| Coast | Pempheridae | Bullseye | *Pempheris* spp | Pempheridae_Pempheris.spp_37357903 |
| Coast | Percophidae | Duckbills | | Percophidae_37393000 |
| Ocean | Phosichthyidae | Lightfishes | | Phosichthyidae_37106913 |
| Tropical | Pinguipedidae | Grub fish | | Pinguipedidae_37390000 |
| Coast | Platycephalidae | Flathead | other | Platycephalidae_other_37296000 |
| Estuary/coast | Platycephalidae | Flathead | *Platycephalus* fuscus | Platycephalidae_Platycephalus.fuscus_37296004 |
| Coast | Plesiopidae | Prettyfins | | Plesiopidae_37316000 |
| Estuary/coast | Pleuronectidae | Flounder | *Rhombosolea* spp | Pleuronectidae_Rhombosolea.spp_37461912 |
| Coast | Pomacentridae | Damselfish | | Pomacentridae_37372000 |
| Estuary/coast | Pomatomidae | Tailor | *Pomatomus saltatrix* | Pomatomidae_Pomatomus.saltatrix_37334002 |
| Coast | Priacanthidae | Bigeyes | | Priacanthidae_37326000 |
| Tropical | Pseudochromidae | Dottyback | | Pseudochromidae_37313000 |
| Estuary/coast | Rhombosoleidae | Flounder | *Ammotretis* spp | Rhombosoleidae_Ammotretis.spp_37461900 |
| Tropical | Samaridae | Flatfish | | Samaridae_37461000.2 |
| Tropical | Scaridae | Parrotfish | | Scaridae_37386000 |
| Tropical | Schindleriidae | Floater | *Schindleria* spp | Schindleriidae_Schindleria.spp_37424901 |
| Estuary/coast | Sciaenidae | Teraglin | *Atractoscion aequidens* | Sciaenidae_Atractoscion.aequidens_37354020 |
| Estuary/coast | Sciaenidae | Mulloway | *Agyrosomus japonicus* | Sciaenidae_Agyrosomus.japonicus_37354001 |
| Coast/pelagic | Scomberesocidae | Saury | *Scomberesox saurus* | Scomberesocidae_Scomberesox.saurus_37236001 |
| Coast/pelagic | Scombridae | Blue mackerel | *Scomber australasicus* | Scombridae_Scomber.australasicus_37441001 |
| Tropical | Scombridae | Tuna | *Auxis* spp | Scombridae_Auxis.spp_37441915 |
| Tropical | Scombridae | Tuna | other | Scombridae_other_37441000 |

| Category | Family-NIMO | Common-NIMO | Species-NIMO | NIMO_NAME |
|---|---|---|---|---|
| Ocean | Scopelarchidae | Pearleyes | | Scopelarchidae_37131000 |
| Coast/slope | Scorpaenidae | Ocean perch | *Helicolenus* spp | Scorpaenidae_Helicolenus.spp_37287918 |
| Coast | Scorpaenidae | Gurnard perch | *Neosebastes* spp | Scorpaenidae_Neosebastes.spp_37287927 |
| Estuary | Scorpaenidae | Cobbler | *Gymnapistes marmoratus* | Scorpaenidae_Gymnapistes.marmoratus_37287018 |
| Coast | Scorpaenidae | | other | Scorpaenidae_other_37287000 |
| Estuary/coast | Scorpaenidae | Fortescue | *Centropogon australis* | Scorpaenidae_Centropogon.australis_37287048 |
| Coast | Scorpididae | Sweep | *Scorpis* spp | Scorpididae_Scorpis.spp_37361906 |
| Coast | Serraninae | Wirrahs | *Acanthistius* spp | Serraninae_Acanthistius.spp_37311912 |
| Tropical | Siganidae | Rabbitfish | *Siganus* spp | Siganidae_Siganus.spp_37438902 |
| Coast | Sillaginidae | Stout whiting | *Sillago robusta* | Sillaginidae_Sillago.robusta_37330005 |
| Coast | Sillaginidae | Eastern school whiting | *Sillago flindersi* | Sillaginidae_Sillago.flindersi_37330014 |
| Coast | Sillaginidae | Southern school whiting | *Sillago bassensis* | Sillaginidae_Sillago.bassensis_37330002 |
| Estuary/coast | Sillaginidae | King George whiting | *Sillaginodes punctatus* | Sillaginidae_Sillaginodes.punctatus_37330001 |
| Coast | Sillaginidae | Whiting | other | Sillaginidae_other_37330000 |
| Estuary/coast | Sillaginidae | Sand whiting | *Sillago ciliata* | Sillaginidae_Sillago.ciliata_37330010 |
| Coast | Soleidae | Sole | | Soleidae_37462000 |
| Estuary/coast | Sparidae | Snapper | *Chrysophrys auratus* | Sparidae_Chrysophrys.auratus_37353001 |
| Estuary/coast | Sparidae | Yellowfin bream | *Acanthopagrus australis* | Sparidae_Acanthopagrus.australis_37353004 |

| Category | Family-NIMO | Common-NIMO | Species-NIMO | NIMO_NAME |
|---|---|---|---|---|
| Estuary/coast | Sparidae | Tarwhine | *Rhabdosargus sarba* | Sparidae_Rhabdosargus.sarba_37353013 |
| Tropical | Sphyraenidae | Barracudas | *Sphyraena* spp | Sphyraenidae_Sphyraena.spp_37382901 |
| Ocean | Sternoptychidae | Hatchet fish | | Sternoptychidae_37107000 |
| Estuary | Syngnathidae | Pipefish | *Stigmatopora nigra* | Syngnathidae_Stigmatopora.nigra_37282018 |
| Estuary | Syngnathidae | Pipefish | other | Syngnathidae_other_37282000 |
| Tropical | Synodontidae | Lizardfish | | Synodontidae_37118000 |
| Estuary/coast | Terapontidae | Trumpeter | *Pelates* spp | Terapontidae_Pelates.spp_37321908 |
| Coast/slope | Tetragonuridae | Squaretail | *Tetragonurus* spp | Tetragonuridae_Tetragonurus.spp_37449901 |
| Estuary/coast | Tetraodontidae | Toadfish | | Tetraodontidae_37467000 |
| Coast | Trachichthyidae | Roughy | *Aulotrachichthys* spp | Trachichthyidae_Aulotrachichthys.spp_37255901 |
| Coast | Trachichthyidae | Roughy | other | Trachichthyidae_other_37255000 |
| Coast/slope | Trichiuridae | Frostfish | *Lepidopus caudatus* | Trichiuridae_Lepidopus.caudatus_37440002 |
| Coast | Trichonotidae | Sanddivers | *Trichonotus* spp | Trichonotidae_Trichonotus.spp_37394901 |
| Coast | Triglidae | Gurnard | *Lepidotrigla papilio* | Triglidae_Lepidotrigla.papilio_37288002 |
| Coast | Triglidae | Gurnard | *Lepidotrigla* spp | Triglidae_Lepidotrigla.spp_37288901 |
| Coast | Triglidae | Gurnard | other | Triglidae_other_37288000 |
| Estuary/coast | Tripterygiidae | Triplefins | | Tripterygiidae_37415000 |
| Coast | Uranoscopidae | Stargazers | | Uranoscopidae_37400000 |
| Tropical | Xiphiidae | Swordfish | *Xiphias gladius* | Xiphiidae_Xiphias.gladius_37442001 |
| Coast/slope | Zeidae | Dory | | Zeidae_37264000 |

**Table S3** Summary of Parameter estimates from the larval fish model. Parameter estimates show the median estimate and 95% credible intervals. If the credible interval does not include zero then it can be considered as an important parameter.

| Parameter | Estimate (95% CI) |
|---|---|
| Upwelling Favourable Winds | -3.52 (-7.65, 0.60) |
| Upwelling Favourable Winds (quadratic) | -7.16 (-10.87, -3.23) |
| Downwelling Favourable Winds | 7.66 (2.43, 12.81) |
| Downwelling Favourable Winds (quadratic) | -1.68 (-5.57, 2.28) |
| Distance to Coast | 0.37 (0.07, 0.68) |
| Upwelling Favourable Winds * Distance to Coast | 6.41 (-0.05, 12.96) |
| Upwelling Favourable Winds (quadratic)* Distance to Coast | 8.23 (0.35, 12.96) |
| Downwelling Favourable Winds * Distance to Coast | -9.08 (-17.46, -1.06) |
| Downwelling Favourable Winds (quadratic)* Distance to Coast | 12.25 (3.23, 21.56) |
| Upwelling Favourable Winds * Downwelling Favourable Winds | 0.16 (-0.02, 0.33) |
| Upwelling Favourable Winds * Downwelling Favourable Winds * Distance to Coast | -0.49 (-0.85, -0.12) |

**Table S4** Summary of Parameter estimates from the CPUE model. Parameter estimates show the median estimate and 95% credible intervals. If the credible interval does not include zero then it can be considered as an important parameter.

| Parameter | Estimate (95% CI) |
|---|---|
| Upwelling Favourable Winds | -161.80 (-303.74, -21.82) |
| Upwelling Favourable Winds (quadratic) | -36.56 (-170.8, 97.8) |
| Downwelling Favourable Winds | -95.38 (-236.77, 44.67) |
| Downwelling Favourable Winds (quadratic) | 21.90 (-134.03, 180.27) |
| Estuary Type: Barrier River | -2.21 (-51.67, 44.87) |
| Estuary Type: Drowned River Valley | 1.61 (-50.59, 51.99) |
| Drought Months | 7.20 (3.67, 10.72) |
| Species: Flathead | -4.66 (-33.49, 23.30) |
| Species: Mullet | 205.48 (101.95, 299.1) |
| Species: Whiting | -38.62 (-68.74, -7.49) |
| Upwelling Favourable Winds * Downwelling Favourable Winds | 3.43 (-8.46, 15.23) |
| Estuary Type: Barrier River: Drought Months | -8.70 (-13.32, -4.16) |
| Estuary Type: Drowned River Valley: Drought Months | -6.69 (-12.02, -1.24) |
